# A holistic survey of small mammal diversity across an iconic Madrean Sky Island (Santa Catalina Mountains, Arizona, USA)

**DOI:** 10.64898/2026.03.15.711934

**Authors:** Dakota M. Rowsey, Stephanie M. Smith, Luisa J. Zamora Chavez, Damien C. Rivera, Savage C. Hess, Matthew F. Jones, Melanie E. Bucci, Shahrzad Mohammadian, Jesse M. Alston, Justin R. Baez, Karla L. Vargas, Nathan S. Upham

## Abstract

The Santa Catalina Mountains are an iconic member of the Madrean Sky Islands, rising above Tucson, Arizona, USA, where the Catalina Highway connects Sonoran desertscrub to stands of conifer forest nearly 2,800 meters in elevation. As one of the ∼54 forested mountain areas in this system, the Santa Catalinas host unique biotic communities relative to the surrounding lowlands. However, most of these sky islands lack the surveys of resident small mammals (either historical or recent) needed for studying biodiversity in the context of changing climate and habitat use. From 2021 to 2023, we surveyed 10 localities on the north and south slopes of the Santa Catalina Mountains using holistic sampling methods to document terrestrial small mammal diversity and preserve multiple tissue types. Here we summarize these new collections relative to previous voucher specimens and human observations, identifying gaps for future work to address. Our survey recorded the presence of 15 species, preserved 150 voucher specimens paired with a suite of flash-frozen tissues, and non-lethally sampled another 219 individuals (ear tissue, feces, ectoparasites, and measurements) to provide populational data from sites where vouchering occurred. Despite the road accessibility and long history of sampling in the Santa Catalina Mountains, our surveys extended the known elevational range for 8 species, including the first known specimen of *Reithrodontomys fulvescens* from the area. Our use of a transect-based survey design, which maximizes species diversity across biotic communities, paired with holistic specimen preservation techniques, provides a model for surveying patterns of population genetic and parasite sharing relationships across other Madrean Sky Islands, bridging a ∼40 year lull in specimen preservation while adding new data dimensions that promote integrative studies of small mammal biodiversity. With more complete sampling, other mountains will offer promising replicates for studying eco-evolutionary impacts of the region’s episodic habitat connectivity.

**Teaser text:** Surveying the terrestrial small mammals of the Santa Catalina Mountains, part of the Madrean Sky Islands, we analyze modern occurrences relative to previous records and demonstrate the potential value of holistically surveying sky island small mammals.

## Introduction

Isolated mountains and mountain ranges surrounded by a less permeable habitat matrix are often referred to as “sky islands” (a special type of “habitat island”; Warshall 1995; Matthews 2021). Sky islands can occur in isolation (called “inselbergs”; Porembski and Barthlott 2012) or multiply, with the latter systems of adjacent sky islands valued as natural laboratories for studying dynamics of speciation, gene flow, local extinction, niche partitioning, exchange of symbionts and parasites, and other ecological processes, particularly as relates to historical environmental changes (Holzmann *et al*. 2023; Love *et al*. 2023; Liu *et al*. 2024; Suzuki *et al*. 2024; see also the other studies in this special feature). The Madrean Sky Island Archipelago of the southwestern United States and northern Mexico is among the best known of the stepping-stone sky island systems globally (Warshall 1995). Many sky islands in the Madrean system exhibit steep relief and distinct elevational bands of different plant assemblages, rising from Sonoran and/or Chihuahuan desertscrub to mixed coniferous forest characteristic of other high peaks in the Rocky Mountain and Sierra Madre Occidental Cordilleras (Pace and Brown 1982). Sandwiched between these two continental mountain ranges, each with distinct species assemblages, the Madrean Sky Islands form a “biogeographic crossroads” for many species of plants and animals (Warshall 1995; Spector 2002; Yanahan and Moore 2019). However, the habitat complexity both within and among mountains has amplified shortfalls of biodiversity knowledge in this region (Koprowski *et al*. 2005; Deyo *et al*. 2013; Rivera and Upham this volume), raising the importance of continued field studies for understanding and protecting the unique biota of the Madrean Sky Islands.

One of the most famous, accessible, and largest members of the Madrean Sky Islands is the Santa Catalina Mountains in southeastern Arizona. They are a prominent geological feature of the Tucson metropolitan area, rising ∼2 kilometers above the Sonoran Desert to the 2,791-meter peak of Mount Lemmon (Moore *et al*. 2013). The Santa Catalinas, known as *Babad Do’ag* in the O’odham language (pronounced *bob-ott doe-awk*), exhibit the characteristic – often abrupt – transitions from desertscrub to conifer forest as one travels from base to peak. Recent surveys of plants and arthropods have documented extensive biodiversity in the Santa Catalina Mountains, including at least 317 plant, 88 ant, and 69 ground beetle species (Brusca *et al*. 2013; Moore *et al*. 2013; Meyer *et al*. 2015). However, the lack of comparable modern surveys of the small mammal or other vertebrate faunas has thus far prevented a more comprehensive view of sky-island biodiversity from taking shape, especially relative to changing temperature, precipitation, and fire regimes in the region (Yanahan and Moore 2019; Wilder *et al*. 2021; Culhane *et al*. 2022). The Santa Catalinas therefore present a key opportunity to study the diversity of small mammals relative to co-occurring species, both now and compared to the historical baseline of preserved voucher specimens in the region. Of particular interest is how biotas have responded to increases in temperature, the duration and severity of drought, the frequency and severity of wildfires, and anthropogenic disturbances like vehicle traffic and outdoor recreation (Pearce and Venier 2005; Letnic *et al*. 2013; Culhane *et al*. 2022). The aridity of this region makes it particularly susceptible to climatic changes and encroaching urbanization (Flesch *et al*. 2010; Yang *et al*. 2016), but baseline diversity data must first be established in order to investigate biotic responses to these changes.

The Santa Catalina Mountains were an early target for surveys by various naturalists and researchers, likely due to the mountains’ large size, proximity to Tucson, and accessibility via the Catalina Highway. William E. D. Scott, an early surveyor, collected over 150 voucher specimens, mostly birds, from south slope forests (Scott 1885a, 1885b); his specimens are, to our knowledge, the first published series of zoological specimens preserved from the Santa Catalina Mountains. Lange (1960) was the first to systematically review mammals from the Santa Catalinas, primarily summarizing the activities of several pioneering collectors rather than conducting original surveys (e.g., Allen 1895; Mearns 1907; Doutt 1934; Dice and Blossom 1937). Lange (1960) is still the most recent treatise on the topic. He documented a total of 73 mammal species and subspecies and another 19 suspected or extirpated species in the Santa Catalina Mountains. Among those, native small mammals constituted 58 species or subspecies in the orders Rodentia (29), Chiroptera (17), and Eulipotyphla (2; see Supplementary Data SD1). However, 95% of all small mammal specimens preserved from the Santa Catalina Mountains were collected prior to 1980 (Fig. 1). Now fully six decades since Lange’s treatise, key questions remain regarding both the basic biodiversity of small mammals in the Santa Catalina Mountains and more applied topics of how those species have responded to ongoing environmental changes. Here we endeavor to tackle the first challenge — which species of terrestrial small mammals are present along the Santa Catalinas elevational gradient?

**Fig. 1.**
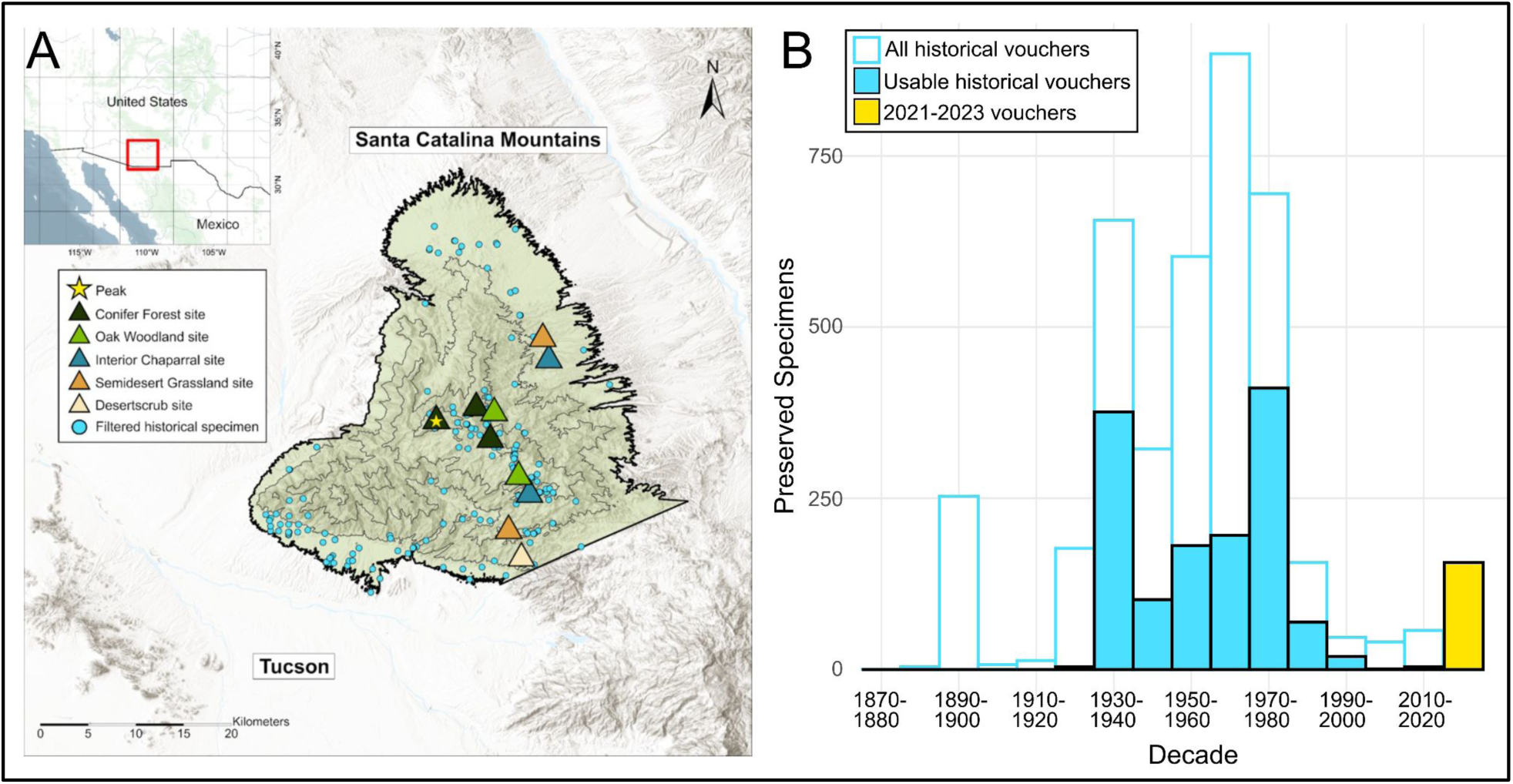
Historical collections relative to our recent survey. **(A)** Map of the study area in the Santa Catalina Mountains, southeastern Arizona, with boundary delineated by the green polygon and topographic contour lines illustrating elevation and terrain variation. Triangles indicate sites sampled during our 2021-2023 survey and are colored according to biotic community. Blue circles represent terrestrial small mammal (Rodentia, Eulipotyphla) specimens collected prior to 2021. The summit of the mountain is denoted by a star. **(B)** Histogram of voucher specimens collected by decade prior to 2021 (white and blue) and in the present study (yellow). Raw records were filtered as described in the Methods. Figure excludes 15 records collected prior to 1850 that were filtered from the final dataset, as well as 233 human observations included in the final analysis but omitted from this figure.

Our study sought to accomplish three goals. First, we aimed to build a modern inventory of small mammals in the Santa Catalina Mountains for comparison to historical data, adding new specimens using a transect-based trapping protocol designed to maximize species diversity across an elevational gradient (Read *et al*. 1988; Bowman *et al*. 2001; Pearson and Ruggiero 2003). Koprowski et al. (2005: 412) highlighted the ‘dearth’ of recent mammal survey data from the Santa Catalinas and other Madrean Sky Islands, but few advances have subsequently been made in the 20 years since that publication. Second, we sought to establish historical context for these new survey results by summarizing all previous records of small mammal vouchers and human observations on these mountains. Our team subsampled 10 of the 66 sites from Moore et al.’s (2013) survey of plants and ground-dwelling arthropods to ensure that biotic communities were evenly sampled for terrestrial small mammals, which also allowed historical records to be interpreted relative to these biotic communities. Third, we aimed to holistically preserve voucher specimens and tissues from our survey to enable future genetic analyses of small mammals and their symbionts across the Madrean Sky Islands — a critical need given the sparse availability of such samples in public databases (see Rivera and Upham, this volume). Holistic preservation methods involve the flash-freezing of multiple tissue types to enable genomic, transcriptomic, and metagenomic analyses (e.g., involving hosts and their viruses, bacteria, or fungi); the ethanol and/or skeletal preservation of whole-organism vouchers to enable in-depth anatomical studies; the sampling of blood (e.g., sera, smears), ectoparasites (e.g., fleas, ticks, mites, botfly larvae), and endoparasites (e.g., cestodes, nematodes, helminths) for a variety of ecological studies; the recording of associated animal measurements and photos; and other specimen-derived data (Galbreath *et al*. 2019; Thompson *et al*. 2021; Timm *et al*. 2021, 2025).

Our transect-based sampling strategy, coupled with presence-only data from historical records, allowed us to make two predictions. First, relative to the species diversity detected per site and across elevations, we expected the diversity of terrestrial small mammals to mirror diversity patterns found in other taxa of the Santa Catalina Mountains. Specifically, we expected the highest mammal diversity to be in the semi-desert grasslands, a midslope region spanning 1330-1645 m where plant and arthropod diversity was shown to peak during surveys in 2011 (Brusca *et al*. 2013; Moore *et al*. 2013) consistent with the mid-domain effect of montane diversity (Colwell and Lees 2000; McCain 2005). Second, relative to the elevational range of each detected small mammal species in the Santa Catalinas, we expected our additional field sampling to yield bidirectional range extensions (i.e., extralimital elevational occurrences) due to statistical chance alone given the sparsity of historical small mammal sampling (∼2 specimens / km^2^; Rivera and Upham, this volume). Alternatively, we expected consistent range extensions up or down slope if underlying environmental drivers of animal movement were the cause (i.e., actual range expansions); however, detailed grid-based sampling for species occupancy would be required to confirm such suspicions (Koprowski *et al*. 2005; Frey 2009). Ultimately, our survey lays the initial groundwork for a broader research program to better understand the past, present, and future of small mammal biodiversity of the Madrean Sky Islands.

## Methods

### Environmental setting

The geological and environmental history of the Santa Catalina Mountains helps contextualize their remarkable biological diversity. The Catalina Fault was formed starting ∼35 million years ago (Ma), during the latest Eocene, with one of the first bouts of extension of the North American plate, part of larger processes of extension across western North America that formed what is now called the Basin and Range Province (Eaton 1982, Coblentz 2005). Deformation of underlying and overlying granites caused bedrock to migrate toward the surface, eventually producing the Catalina Metamorphic Core Complex (Bezy 2016). Extension halted in the Santa Catalina region ∼5 Ma, and erosion became the primary factor shaping the range. Today, granites and gneisses, being more resistant than other rocks, form most of the dominant surface features of the Santa Catalina Mountains (Bezy 2016).

The Santa Catalinas host a steep environmental gradient, with mean temperature decreasing by ∼12 °C and annual precipitation increasing by ∼50 cm from base to peak (Whittaker and Niering 1975), which has produced distinctive biotic communities that roughly track elevation (Brown and Lowe 1980; Brown 1982:198). We use the definitions provided by Brown (1982), who described the biotic communities of increasing elevation in the Santa Catalina Mountains as follows: (i) Sonoran Desertscrub, Upland Division (which we abbreviate as Desertscrub, from the base of the mountain’s southern aspect at ∼830 to 1,500 m); (ii) Semidesert Grassland (∼830 to 1,820 m); (iii) Interior Chaparral (∼932 to 1,857 m); (iv) Madrean Evergreen Woodland (sometimes referred to as Madrean Oak Woodland, and which we abbreviate as Oak Woodland, ∼875 to 2,550 m); and (v) Petran Montane Conifer Forest (which we abbreviate as Conifer Forest, ∼1,600 to the mountain’s peak at ∼2,800 m; (NASA/METI/AIST/Japan Spacesystems and ASTER Science Team 2019). As their names suggest, these biotic communities are defined primarily by their dominant plants. Desertscrub is dominated by plants adapted to arid conditions and highly seasonal and ephemeral availability of free water, such as yellow palo verde (*Parkinsonia microphylla*), cholla (*Cylindropuntia* spp.), mesquite (*Neltuma* [*Prosopis*] spp.), and the characteristic saguaro cactus (*Carnegiea gigantea*). Semidesert Grassland is similar in some respects to the biotic communities of the Chihuahuan desert, and characterized by grasses and shrubs including sideoats grama (*Bouteloa curtipendula*), cane beardgrass (*Bothriochloa barbinodis*), and the introduced Lehmann lovegrass (*Eragrostis lehmanniana*), as well as agaves (*Agave* spp.) and other succulents (Judd 1962). Interior Chaparral forms patchy clusters based partially on elevation, but is influenced by elevation-independent aspect and soil moisture availability more than other biotic communities, with dominant plant species including shrub oak (*Quercus turbinella*), manzanitas (*Arctostaphylos* spp.), and desert ceanothus (*Ceanothus pauciflorus*). Oak Woodland sees the expansion of small trees into an open-canopy wooded habitat with a prevalence of species such as Arizona pine (*Pinus arizonica*), alligator-bark juniper (*Juniperus deppeana*), and Emory oak (*Quercus emoryi*). Finally, Conifer Forest is restricted to areas near the peak of the mountains, forming denser, closed-canopy forest characterized by species such as Arizona pine (*Pinus arizonica*), southwestern white pine (*Pinus strobiformis*), and Douglas-fir (*Pseudotsuga menziesii*), occasionally accompanied by clusters of aspens (*Populus tremuloides*). The boundaries between these communities are often stark, with relatively narrow ecotones between them, except in areas highly disturbed by human activity or wildfire, which can allow plant species from lower-elevation adjacent biotic communities to penetrate into higher elevations (Bennett *et al*. 2004; Brusca and Moore 2013; Moore *et al*. 2013; Yanahan and Moore 2019). Changes in biotic communities caused by human disturbance have occurred alongside elevational shifts caused by climate change, leading to a contraction of montane woodland and forest (Brusca *et al*. 2013) and an expansion of grasslands (Wilder *et al*. 2021).

### Field methods

We conducted field surveys from November 2021 to July 2023 to sample terrestrial small mammals in each of the major biotic communities along the elevational gradient of the Santa Catalina Mountains (Fig. 1A, 2A). We sampled at 10 sites—5 along the south aspect of the mountain, 4 along the north aspect, and 1 at the peak—corresponding to the 5 major biotic communities of plants and animals that traverse the mountain range (Brown 1982; Moore et al. 2013; Table 1): (i) Sonoran Desertscrub (south aspect only); (ii) Semidesert Grassland (north and south); (iii) Interior Chaparral (north and south); (iv) Oak Woodland (north and south); and (v) Conifer Forest (north, south, and peak). Two of the sites we sampled – the north slope Oak Woodland (mean elevation 2,075 m) and the peak Conifer Forest (mean elevation 2,759 m) – were burned during the 2020 Bighorn Fire, which burned 486 km^2^ of the study area between 5 June and 23 July 2020 (i.e., ∼57% of the mountain’s ∼850 km^2^ total area, as defined below; Wilder et al. 2021). We determined site burn status in the field through visual observation and confirmed these observations with data from the National Interagency Fire Center (NIFC Authoritative Content 2023).

**Fig. 2.**
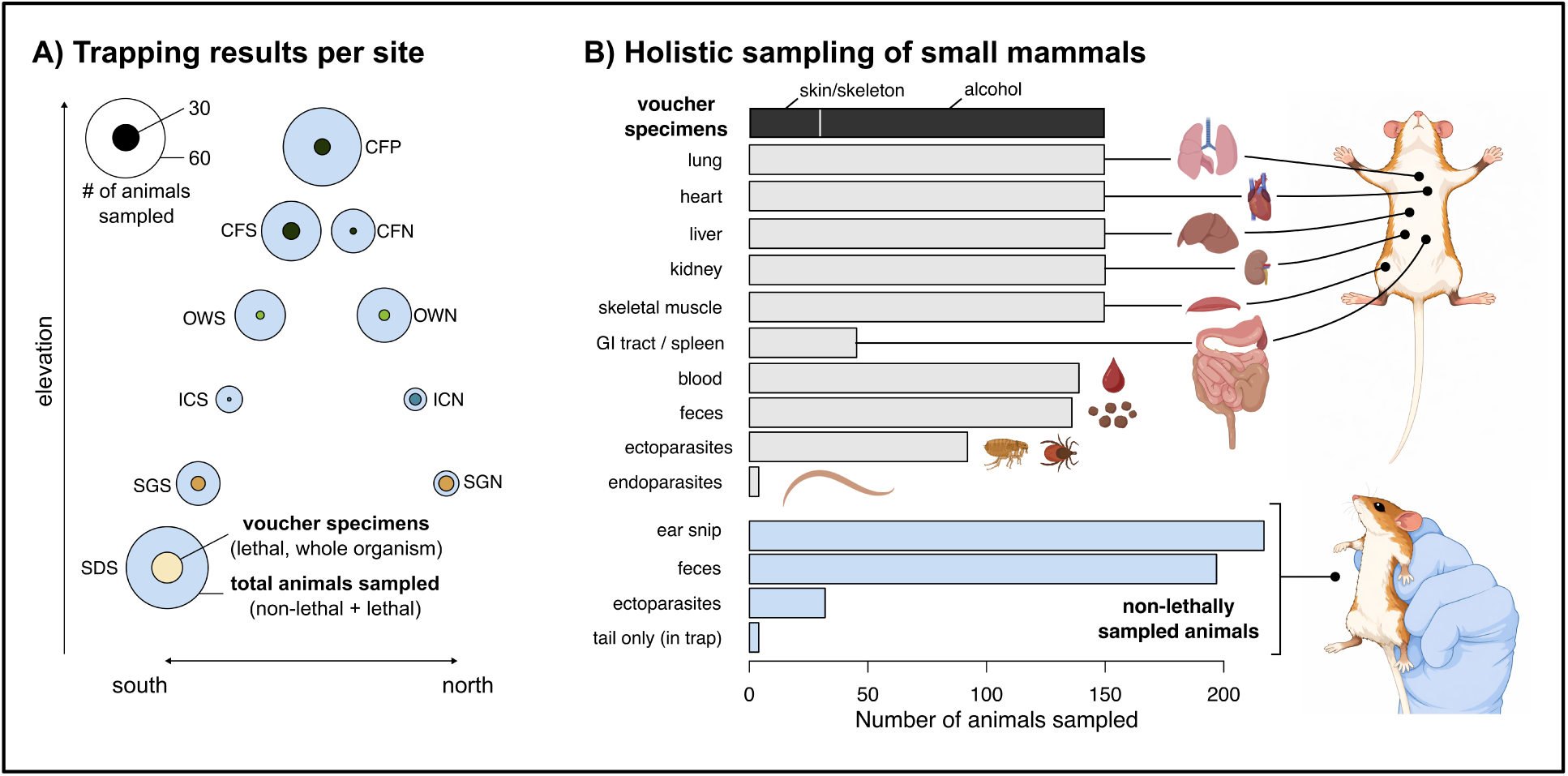
Overview of our 2021-2023 survey of the Santa Catalina Mountains. **(A)** Trapping results for each of the 10 sites sampled across elevations on the north and south mountain slopes. Site abbreviations and elevations are given in Table 1. Note that higher elevations exclude desertscrub habitat on the mountain’s north side. **(B)** Summary of the animals sampled for each tissue type, both for voucher specimens (lethal, n=150) and human observations (non-lethal, n=219), with all tissues preserved via flash-freezing on liquid nitrogen or dry ice and storage at - 80C. Note that not all sampled animals had available feces or parasites, some animals were missed for blood samples, and gastrointestinal (GI) tracts were only sampled for the first vouchered animal/species/site. All animals (n=364) have external measurement data and/or photographs of ventral, dorsal, and lateral views. Images are from BioRender as modified in some cases using ChatGPT v5.2.

**Table 1.** Details of the 10 sites sampled across the Santa Catalina Mountains from 2021-2023. Latitude and longitude of the site were computed from the centroid of sampled specimens around each site.

| Site | Biotic Community | Slope | Latitude | Longitude | Elevation Range (m) |
| --- | --- | --- | --- | --- | --- |
| SDS | Desertscrub | S | 32.31322438 | -110.7123025 | 1146 - 1214 |
| SGS | Semidesert Grassland | S | 32.33744 | -110.71924 | 1460 - 1538 |
| ICS | Interior Chaparral | S | 32.37734268 | -110.6857492 | 1884 - 1929 |
| OWS | Oak Woodland | S | 32.39200448 | -110.7047277 | 2096 - 2163 |
| CFS | Conifer Forest | S | 32.42276744 | -110.7317724 | 2363 - 2431 |
| CFP | Conifer Forest | Peak | 32.4412 | -110.78434 | 2761 - 2790 |
| CFN | Conifer Forest | N | 32.45156243 | -110.7467846 | 2195 - 2391 |
| OWN | Oak Woodland | N | 32.45021099 | -110.7373919 | 2055 - 2095 |
| ICN | Interior Chaparral | N | 32.50020822 | -110.6825961 | 1315 - 1342 |
| SGN | Semidesert Grassland | N | 32.51622 | -110.68707 | 1348 - 1368 |

For each of the 10 sites, we sampled mammals for a total of 400 trapnights (4,000 trapnights total for the survey) by setting 200 Sherman folding live-traps (3 x 3.5 x 9”) in transects of 3-5 lines, each with 40-80 traps, and opening traps at dusk for 2 consecutive nights. Traps were baited with rolled oats and sterilized bird seed, checked at dawn, and closed during peak daylight heat (between ca. 09:00 and 16:00). Traps were placed in pairs ∼2 m apart from each other at trapping stations, with each station ∼10 m apart, in areas of continuous habitat conducive to sampling small mammals (e.g., shaded, concealed near rocks or vegetation, near burrows, and/or nearby mammal signs of prints, feces, or tail drag marks).

Up to 3 individuals per species per site (the maximum allowed under our scientific collecting permits) were euthanized with vapor-phase isoflurane and prepared as voucher specimens. These vouchers were preserved as whole fluid-preserved specimens, initially fixed in formalin then preserved in ethanol; similarly preserved specimens with skulls extracted; or stuffed study skins and skeletons. All vouchers and tissues were deposited in the Arizona State University Mammalogy Collection (ASUMAC) at the ASU Natural History Collection (ASUNHC). We also vouchered all specimens found deceased in our traps as salvage (< 2% of trapnights overall). We sampled a standard array of tissues from all voucher specimens: skeletal muscle, liver, kidney, heart, lung, Nobuto blood smears, feces, and ecto/endo-parasites when available (Fig. 2B). These samples were flash-frozen on dry ice or liquid nitrogen in the field and transferred to the ASU Vertebrate Tissue Collection for archival storage (ASUVTC). We also non-lethally sampled animals that exceeded the maximum lethal take of our permits by collecting ear clips, fecal samples, ectoparasites, external measurement data, and/or photographs of ventral and lateral views for future validation of IDs before releasing the animals alive at the site of capture, with individuals being identified based on external characteristics. We conducted all field sampling procedures in accordance with Arizona Game and Fish Department regulations (AZGFD Scientific Activity License numbers SP407147, SP623885, SP939945), United States Forest Service regulations (Coronado National Forest Special Use Permit numbers SAN2139, SAN2218, SAN2329), and ASU Institutional Animal Care and Use Committee protocols (21-1849R), as well as guidelines for the use of live mammals in research by the American Society of Mammalogists (Sikes *et al*. 2016).

### Study area delimitation

To delineate the study area for comparison with historical biodiversity occurrence data, we used ArcGIS Pro (ESRI 2024) and the ASTER Global Digital Elevation Model (DEM; NASA/METI/AIST/Japan Spacesystems and U.S./Japan ASTER Science Team 2019), applying two elevation-based thresholds relative to the summit of Mount Lemmon (32.4422° N, 110.7886° W). First, on the northern and eastern flanks of the Santa Catalinas, we set a minimum elevation threshold of 1,220 m to reflect where the northern slopes begin to rise distinctly from the San Pedro Valley (Whittaker and Niering 1968). Second, on the southern flank, we set a lower threshold of 840 m to distinguish the transition zone where the mountain range gives way to the Catalina Foothills and Tanque Verde areas (U.S. Department of Agriculture, Forest Service 2025a,b). Using those thresholds, we then separated the Santa Catalinas from the adjacent Rincon Mountains by visually inspecting the DEM layers of these mountains to determine a topographic dividing point. We identified this boundary at Agua Caliente and Bullock Canyons, rather than the commonly cited Redington Pass (Sky Island Alliance 2008), which has greater elevational affinity with the Rincon Mountains.

### Historical data acquisition and quality control

To establish a historical record of mammal occurrences in the Santa Catalina Mountains, we queried the Global Biodiversity Information Facility (GBIF) for both voucher specimen records and iNaturalist research-grade observations (GBIF.org 2024; Telenius 2011). We used a simplified four-vertex polygon for downloading GBIF data (32.70782, -110.99009; 32.18734, - 110.95899; 32.37179, -110.51584; 32.66954, -110.61576; doi: 10.15468/dl.yx4qd8; GBIF.org 2024). We obtained 4,906 mammal records from this initial download on 23 January 2024. We then uploaded these raw records to ArcGIS Pro for filtering to our more complex study-area polygon. In addition to these data, we also downloaded records of all mammal specimens collected from Pima and Pinal counties held by the Smithsonian National Museum of Natural History (USNM; *n* = 940) and the American Museum of Natural History (AMNH; *n* = 28) directly from these institutions’ data portals since these datasets are not available on GBIF. We refined the records from USNM to include only the georeferenceable specimens from within the study-area polygon, for a total of 28 records from the USNM (56 total from both sources; note: our search returned many USNM records from US-Mexico border surveys, which were conducted well south and west of our study area and therefore not relevant to our study).

We examined specimen records from the GBIF download and removed unreliable records based on errors such as imprecise georeferencing and incorrect specimen identification, problems which are exacerbated when fine spatial resolution is necessary (Zizka et al. 2020). We used OpenRefine v. 3.8.1 (Delpeuch *et al*. 2024) to search our dataset, group individuals by locality, and prune records with locality information too imprecise for the level of precision required by our study (e.g., “10 miles N, 8 miles E Tucson”) or with descriptive localities that did not match occurrence record coordinates. We also manually examined outlying elevational values at the species level, verifying specimen identification and occurrence data with collection staff when possible, and removed records with locality information too vague to calculate elevation (see repeatable workflows in the Github that accompanies this article). The resulting filtered dataset contained 2,511 occurrence records.

We then assigned elevational values to all records using the ASTER DEM elevation based on their coordinate location for comparison to records with elevation provided in the occurrence record itself (i.e., verbatim elevation). We performed a linear regression of DEM elevation versus verbatim elevation for the 1,847 records for which these data were available. We extracted regression residuals and excluded 19 records with residuals that exceeded 3 standard deviations above the mean residual value (298.5 meters). This exclusion threshold was chosen because those records showed no positive correlation between DEM elevation and verbatim elevation (Supplementary Data SD2). We performed a manual inspection of the resulting records to examine any suspicious elevational values or species irrelevant to the study, such as introduced or clearly misidentified taxa (e.g., endemic to regions well outside our study system). Taxonomic updates were made following v2.0 of the Mammal Diversity Database taxonomy (Burgin et al. 2025; Mammal Diversity Database 2025). After removing 22 dubious records from subsequent analyses, we filtered the dataset to include only records of Eulipotyphla and Rodentia for more appropriate comparisons to our field sampling. The historical dataset spanning 1916–2020 included 1,367 voucher specimens and 233 human observations for a total of 1,600 records.

We calculated sampling bias associated with elevation, terrain ruggedness, and distance from the nearest road, using the DEM information described previously and local road data obtained from the U.S. Census Bureau’s TIGERweb database, which we clipped to the spatial extent of the study area polygon (United States Census Bureau 2024). We calculated terrain ruggedness as the mean of the absolute differences between the raster cell in which each specimen was collected and the value of the 8 surrounding cells using the package *terra* v. 1.8-60 (Wilson et al. 2007, Hijmans 2025). We calculated the nearest-neighbor distance between occurrence records and the TIGERweb road data using the R package *nngeo* v. 0.4.8 (Dorman 2024). The observed values of elevation, terrain ruggedness, and distance from roads were compared to values generated from 10,000 simulated sampling locations on the mountain, with significance tested using one-way MANOVA.

### Analytical methods

To visualize the association between small mammal species and biotic communities, we conducted correspondence analysis using the R package *ca* version 0.71.1 (Greenacre et al. 2020). We first created a presence-absence matrix for all relevant taxa across all biotic communities. We then calculated Bray-Curtis distance among assemblages using the vegdist function in package *vegan* version 2.6-4 (Oksanen *et al*. 2022), and executed an ordination using Principal Coordinate Analysis (PCoA) via the function “cmdscale()”. To compare small mammal diversity from our survey with plant diversity previously surveyed at the same sites, we extracted values from Table 6 in Moore et al. (2013) and collated the species richness of comparable biotic communities. We mapped the following biotic community categories for comparison: SDS to “Desertscrub (S)”; SGS to “Oak-Grassland (S)”; ICS to “Chaparral (S)”; OWS to “Pine-Oak Woodland (S)”; CFS to Pine Forest (S)”; CFP to Mixed Conifer Forest (MTN); CFN to “Pine Forest (N)”; OWN to “Oak Woodland (N)”; ICN to “Chaparral (N)”; and SGN to “Desert Grassland (N)”. Two categories were unmapped due to lack of analogy with our sampled sites: “Grazing-Disturbed Grassland (N)” and “Pine-Oak Woodland (N)”. We repeated this procedure for Meyer et al. (2015) S2 data to compare our small mammal diversity values to ground-dwelling arthropod diversity from the same sites, mapping the biotic community categories as follows: SDS to “DS-S”; SGN to “GL-N”; SGS to “GL-S”; ICN to “CH-N”; ICS to “CH-S”; OWN to “PO-N”; OWS to “PO-S”; CFN to “P-N”; CFS to “P-S”; CFP to “MC” (unmapped categories were “DS-N”, “OW-N”, “P-C”, and “MC-S”). All files needed to replicate our data analysis in R are available in the project GitHub repository at https://github.com/rowseydm/sc_mamm_survey.

## Results

### Modern field survey

We collected 150 voucher specimens and 1,651 tissue samples from 369 individual animals, 364 of which were identifiable to species, including both vouchered and nonlethally sampled specimens (Fig. 1, Fig. 2). The digital listing of these records in CVColl are available here: vouchers: https://cvcoll.org/portal/collections/listtabledisplay.php?datasetid=9; tissues: https://cvcoll.org/portal/collections/listtabledisplay.php?datasetid=10. We recorded 15 species in 9 genera from the families Cricetidae (9 spp.), Heteromyidae (3 spp.), Sciuridae (2 spp.), and Soricidae (1 sp.; Table 2). Captures represented an overall mean trap-success rate of 9.2% (range by night: 0.5–25%). Trap success varied among biotic communities, ranging from Interior Chaparral (4.6%) to Desertscrub (16%; Table 2). Our highest species richness was in Semidesert Grassland (9 species), followed by Conifer Forest (7), and then Desertscrub, Interior Chaparral, and Oak Woodland (6 species each; Table 2).

**Table 2.** Summary of species captured by site for the 364 captures identifiable to species. Parenthetic numbers in column headers are the trapnight totals for each biotic community, as grouped across sites on the north and south slope and peak; those for each species indicate relative capture success per trapnight (i.e., trap success). Overall and per-species trap success is given at the bottom relative to 4,000 trapnights in the survey. Trap success is underestimated for species of *Peromyscus*, *Chaetodipus*, and *Neotoma* (and overall) given that species-level identifications required handling that at some sites was infeasible due to high abundances.

|  | **Biotic Community** (number of trapnights) |  |  |  |  |  |
| --- | --- | --- | --- | --- | --- | --- |
| **Species** | Desertscrub<br>(400) | Semidesert<br>Grassland<br>(800) | Interior<br>Chaparral<br>(800) | Oak<br>Woodland<br>(800) | Conifer<br>Forest<br>(1200) | **Totals** |
| **Cricetidae** |  |  |  |  |  |  |
| *Neotoma albigula* | 10 (0.025) | 5 (0.00625) | 0 | 2 (0.0025) | 0 | 17 (0.00425) |
| *Neotoma mexicana* | 0 | 0 | 0 | 2 (0.0025) | 18 (0.015) | 20 (0.005) |
| *Onychomys torridus* | 0 | 1 (0.00125) | 3 (0.0075) | 0 | 0 | 4 (0.001) |
| *Peromyscus boylii* | 0 | 12 (0.0015) | 24 (0.06) | 40 (0.05) | 36 (0.03) | 112 (0.028) |
| *Peromyscus eremicus* | 20 (0.05) | 3 (0.00375) | 2 (0.005) | 0 | 0 | 25 (0.0063) |
| *Peromyscus melanotis* | 0 | 0 | 0 | 0 | 57<br>(0.0475) | 57 (0.0142) |
| *Reithrodontomys*<br>*fulvescens* | 0 | 0 | 0 | 0 | 1<br>(0.00083) | 1 (0.0003) |
| *Reithrodontomys megalotis* | 0 | 0 | 0 | 10 (0.0125) | 7<br>(0.00583) | 17 (0.0043) |
| *Sigmodon ochrognathus* | 2 (0.005) | 1 (0.00125) | 0 | 6 (0.0075) | 0 | 9 (0.0023) |
| **Heteromyidae** |  |  |  |  |  |  |
| *Chaetodipus baileyi* | 13 (0.0325) | 7 (0.00875) | 5 (0.0125) | 0 | 0 | 25 (0.0063) |
| *Chaetodipus intermedius* | 18 (0.045) | 16 (0.02) | 2 (0.005) | 0 | 0 | 36 (0.009) |
| *Chaetodipus penicillatus* | 0 | 5 (0.00625) | 0 | 0 | 0 | 5 (0.0013) |
| **Sciuridae** |  |  |  |  |  |  |
| *Ammospermophilus harrisi* | 1 (0.0025) | 2 (0.0025) | 0 | 0 | 0 | 3 (0.0008) |
| *Neotamias dorsalis* | 0 | 0 | 1 (0.0025) | 17<br>(0.02125) | 14<br>(0.0117) | 32 (0.008) |
| **Soricidae** |  |  |  |  |  |  |
| *Sorex monticolus* | 0 | 0 | 0 | 0 | 1<br>(0.00083) | 1 (0.0003) |
| **Totals** | **64 (0.16)** | **52 (0.065)** | **37**<br>**(0.04625)** | **77**<br>**(0.09625)** | **134**<br>**(0.1117)** | **364 (0.091)** |

No single species was sampled from all biotic communities; however, *Peromyscus boylii* was the most frequently captured species, with 112 individuals sampled across 4 of the 5 communities (all but Desertscrub). *Sorex monticolus* and *Reithrodontomys fulvescens* were the least frequently captured species (1 individual each in Conifer Forest; Table 2). Several species were sampled in multiple communities: *Neotoma albigula* and *Sigmodon ochrognathus* were captured from 1,174 m to 2,094 m, spanning from Desertscrub to burned Oak Woodland; other species were restricted to closed-canopy forest, such as *Reithrodontomys megalotis* (2,064–2,423 m) and *Peromyscus melanotis* (2,378–2,789 m; see Discussion for more information regarding our choice of nomenclature). Heteromyids were only found in low- to mid-elevation Desertscrub and Semidesert Grassland (1,149–1,538 m). We observed the lowest degree of community overlap between Interior Chaparral and Oak Woodland, where only 2 of 10 total species were shared.

We quantified species associations with biotic communities using Principal Coordinate Analysis (PCoA). We found that Dimension 1 of the PCoA explained 59% of dataset variance, whereas Dimension 2 explained 19% (Fig. 3; Supplementary Data SD3). All 4 dimensions were needed to explain 95% of dataset variance. Dimension 1 had a clear signal of elevational plant communities, with Conifer Forest exhibiting the highest scores, Desertscrub exhibiting the lowest scores, and other communities occurring in order from lowest to highest elevations (Fig. 3). Dimension 2 is defined by the difference between Interior Chaparral and Oak Woodland communities (Fig. 3). These two communities share only two species: *Neotamias dorsalis* and *Peromyscus boylii*, meaning that they share the fewest species between pairs of elevationally adjacent biotic communities (Supplementary Data SD3). However, they are farther apart from each other on Dimension 2 than are Desertscrub and Conifer Forest, which share 0 species. In terms of taxa, extremes of Dimension 2 are occupied by *Onychomys torridus* (negative), and *Sigmodon ochrognathus* + *Neotoma albigula* (positive, with identical occurrence patterns). Dimension 3 (not shown) explained 13.95% of dataset variance but did not have a clear signal with regard to taxa or biotic communities.

**Fig. 3.**
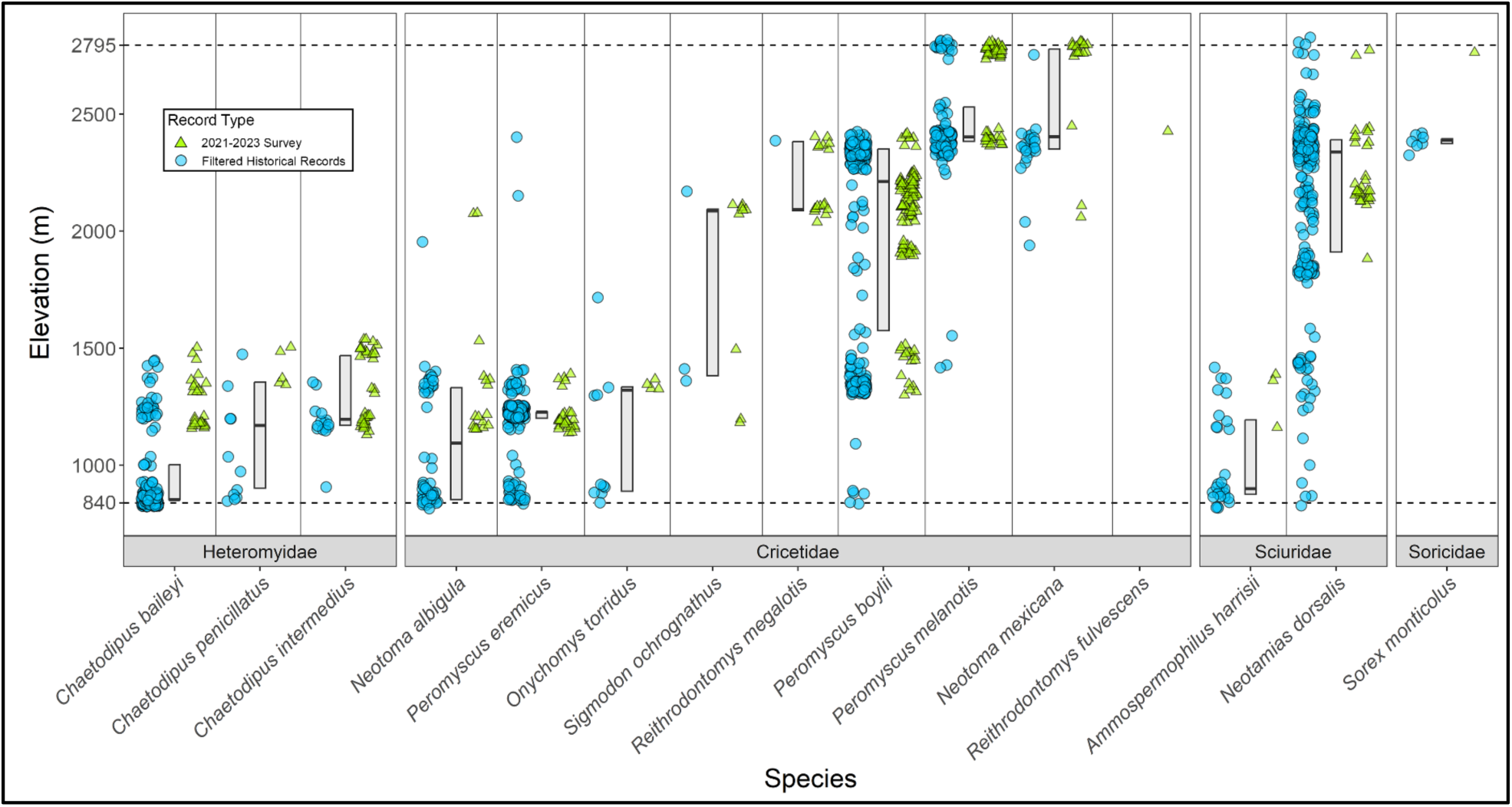
Beeswarm plot of recorded elevations of historical specimens (blue circles) and specimens collected in this survey (green triangles). The boxplots show the interquartile range of the specimens’ elevational distribution and the vertical dashed lines represent the lower (840 m) and upper (2,795 m) boundaries of the mountain. *Peromyscus melanotis* here combines all occurrences labeled *P. maniculatus, P. sonoriensis, P. melanotis,* and *P. leucopus* (see Discussion).

### Modern and historical records: assessment and comparison

We found verifiable historical records of 27 species of terrestrial small mammals in the Santa Catalina Mountains (Table 3). Of those, 19 of 27 species were recorded below 1,000 m, indicating some presence within conservative estimates of the Desertscrub elevational band (Whittaker and Niering 1965). In parallel, 14 species had a maximum elevation of at least 2,200 m, indicating some presence in Conifer Forest, while 18 species reach at least 1,600 m consistent with presence in the forests of the Madrean Sky Islands (Moore et al. 2013; see also Rivera and Upham, this volume). Taking presence of ≥15 historical records as indicative of minimum baseline sampling effort, we find that 6 of the 18 species meeting this criteria appear to be habitat generalists, with elevational ranges ≥1500 m consistent with presence in all biotic communities (*Neotamias dorsalis, Otospermophilus variegatus, Sciurus aberti, Megascapheus bottae, Peromyscus boylii*, and *Peromyscus eremicus*). In contrast, only 3 of those 18 species appear to be Conifer Forest specialists, with >95% of records above 2,000 m (*Peromyscus melanotis, Neotoma mexicana*), and 1 species appears to be restricted below 1,000 m suggesting Desertscrub-Semidesert Grassland specialization (*Sigmodon arizonae*; Table 3).

**Table 3.**
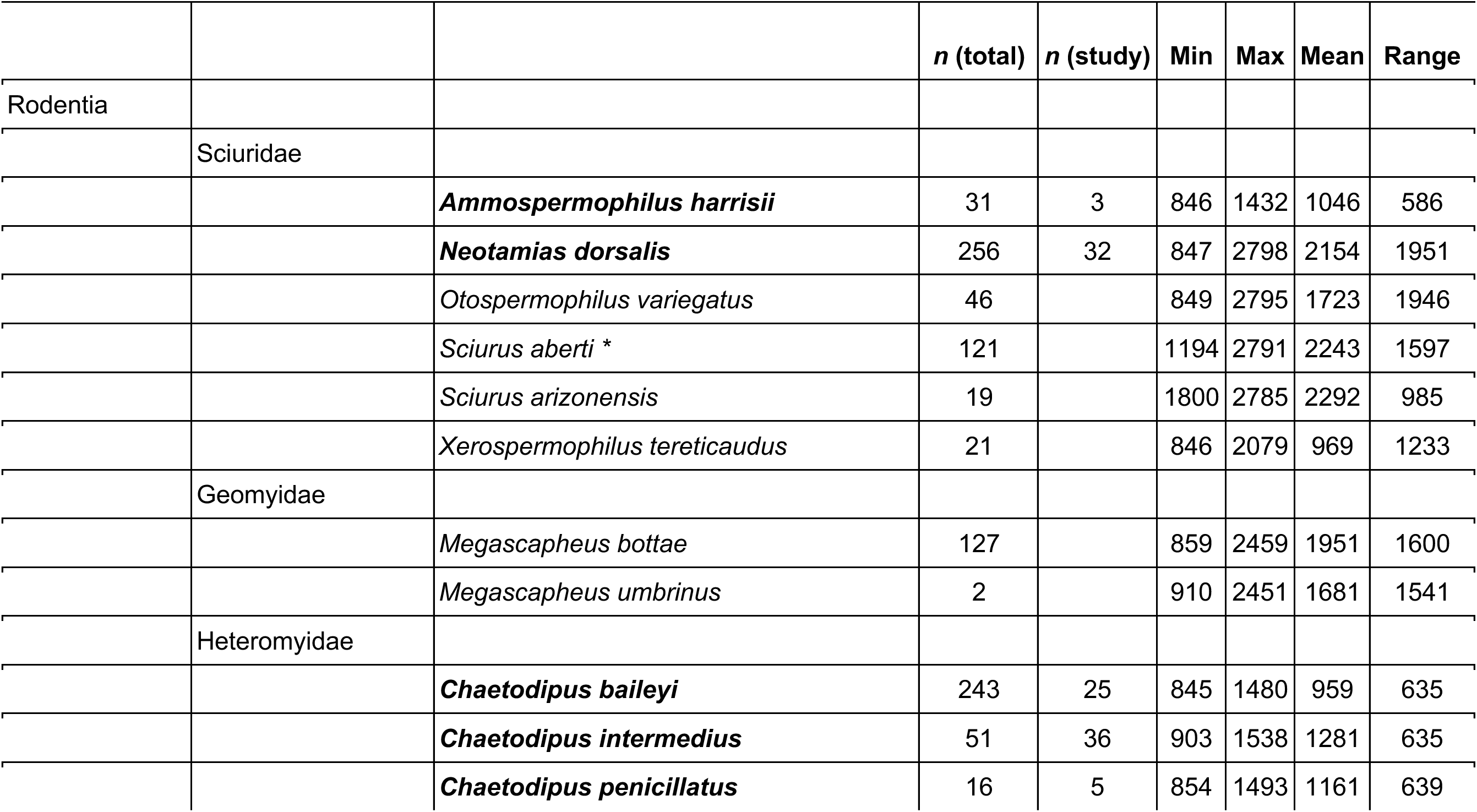
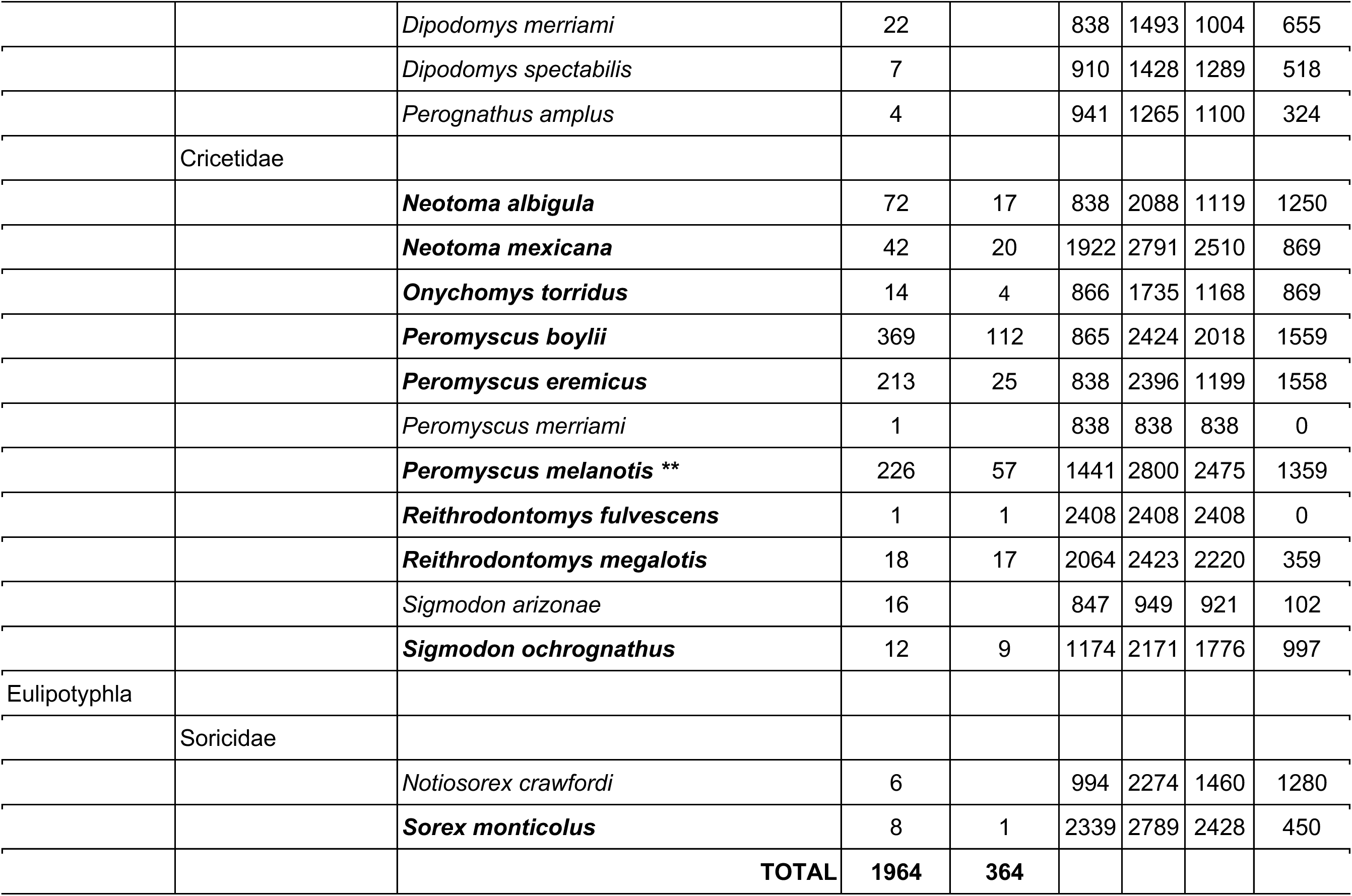
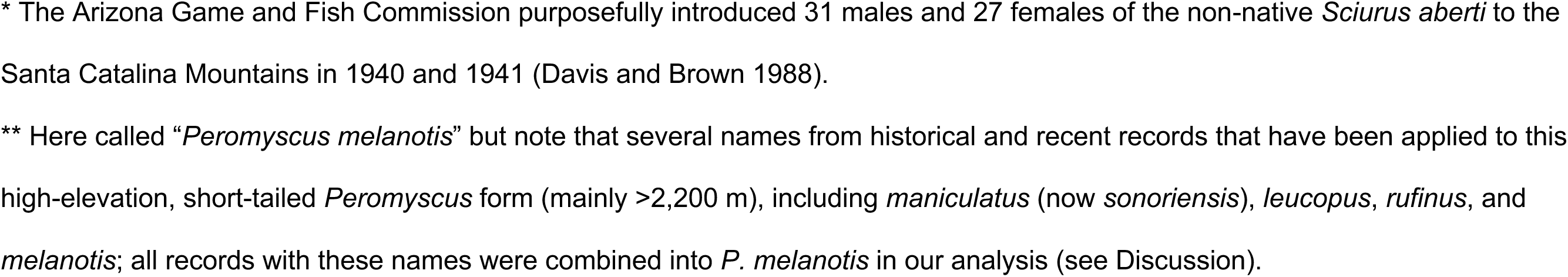
Total (and this study’s) sampling of terrestrial small mammal species by taxon in the Santa Catalina Mountains (*n* includes both voucher specimens and non-lethal observations). The elevational minimum (Min), maximum (Max), mean, and range in meters (m) is shown for each taxon, with the minimum restricted to 840 m (lower limit of the study area). Bolded taxa were captured in the modern survey of this study (2021-2023); 5 non-lethally sampled animals identified to genus are excluded from this tabulation.

|  |  |  | *n* (total) | *n* (study) | Min | Max | Mean | Range |
| --- | --- | --- | --- | --- | --- | --- | --- | --- |
| Rodentia |  |  |  |  |  |  |  |  |
|  | Sciuridae |  |  |  |  |  |  |  |
|  |  | ***Ammospermophilus harrisii*** | 31 | 3 | 846 | 1432 | 1046 | 586 |
|  |  | ***Neotamias dorsalis*** | 256 | 32 | 847 | 2798 | 2154 | 1951 |
|  |  | *Otospermophilus variegatus* | 46 |  | 849 | 2795 | 1723 | 1946 |
|  |  | *Sciurus aberti* * | 121 |  | 1194 | 2791 | 2243 | 1597 |
|  |  | *Sciurus arizonensis* | 19 |  | 1800 | 2785 | 2292 | 985 |
|  |  | *Xerospermophilus tereticaudus* | 21 |  | 846 | 2079 | 969 | 1233 |
|  | Geomyidae |  |  |  |  |  |  |  |
|  |  | *Megascapheus bottae* | 127 |  | 859 | 2459 | 1951 | 1600 |
|  |  | *Megascapheus umbrinus* | 2 |  | 910 | 2451 | 1681 | 1541 |
|  | Heteromyidae |  |  |  |  |  |  |  |
|  |  | ***Chaetodipus baileyi*** | 243 | 25 | 845 | 1480 | 959 | 635 |
|  |  | ***Chaetodipus intermedius*** | 51 | 36 | 903 | 1538 | 1281 | 635 |
|  |  | ***Chaetodipus penicillatus*** | 16 | 5 | 854 | 1493 | 1161 | 639 |
|  |  | *Dipodomys merriami* | 22 |  | 838 | 1493 | 1004 | 655 |
|  |  | *Dipodomys spectabilis* | 7 |  | 910 | 1428 | 1289 | 518 |
|  |  | *Perognathus amplus* | 4 |  | 941 | 1265 | 1100 | 324 |
|  | Cricetidae |  |  |  |  |  |  |  |
|  |  | ***Neotoma albigula*** | 72 | 17 | 838 | 2088 | 1119 | 1250 |
|  |  | ***Neotoma mexicana*** | 42 | 20 | 1922 | 2791 | 2510 | 869 |
|  |  | ***Onychomys torridus*** | 14 | 4 | 866 | 1735 | 1168 | 869 |
|  |  | ***Peromyscus boylii*** | 369 | 112 | 865 | 2424 | 2018 | 1559 |
|  |  | ***Peromyscus eremicus*** | 213 | 25 | 838 | 2396 | 1199 | 1558 |
|  |  | *Peromyscus merriami* | 1 |  | 838 | 838 | 838 | 0 |
|  |  | ***Peromyscus melanotis* \*\*** | 226 | 57 | 1441 | 2800 | 2475 | 1359 |
|  |  | ***Reithrodontomys fulvescens*** | 1 | 1 | 2408 | 2408 | 2408 | 0 |
|  |  | ***Reithrodontomys megalotis*** | 18 | 17 | 2064 | 2423 | 2220 | 359 |
|  |  | *Sigmodon arizonae* | 16 |  | 847 | 949 | 921 | 102 |
|  |  | ***Sigmodon ochrognathus*** | 12 | 9 | 1174 | 2171 | 1776 | 997 |
| Eulipotyphla |  |  |  |  |  |  |  |  |
|  | Soricidae |  |  |  |  |  |  |  |
|  |  | *Notiosorex crawfordi* | 6 |  | 994 | 2274 | 1460 | 1280 |
|  |  | ***Sorex monticolus*** | 8 | 1 | 2339 | 2789 | 2428 | 450 |
|  |  | **TOTAL** | **1964** | **364** |  |  |  |  |

\* The Arizona Game and Fish Commission purposefully introduced 31 males and 27 females of the non-native *Sciurus aberti* to the Santa Catalina Mountains in 1940 and 1941 (Davis and Brown 1988). \*\* Here called "*Peromyscus melanotis*" but note that several names from historical and recent records that have been applied to this high-elevation, short-tailed *Peromyscus* form (mainly >2,200 m), including *maniculatus* (now *sonoriensis*), *leucopus*, *rufinus*, and *melanotis*; all records with these names were combined into *P. melanotis* in our analysis (see Discussion).

Combined historical (pre-2021) and recent (2021-2023) sampling of the Santa Catalina Mountains is biased with respect to elevation, terrain ruggedness index (TRI), and distance from nearest roads (one-way MANOVA: approx. *F* = 1170.3, *P* < 0.001; Fig. 4, Table 5). Sampling is biased toward high elevations (<u>*x*</u>_*obs*_: 1768 m, <u>*x*</u>_*sim*_: 1514 m, *P* < 0.001; Table 5, Fig. 4A), with dense sampling around 2,300 m from historical records and 2,200 m from recent records relative to the amount of area at that elevation (Figure 4A, Supplementary Data SD4A). Additionally, historical sampling is biased toward less rugged areas (<u>*x*</u>_*obs*_: 6.39, <u>*x*</u>_*sim*_: 7.76 m, *P* < 0.001; Table 5), with the maximum observed TRI (22.9) less than half of the maximum simulated TRI (48.9; Figure 4B). Finally, sampling of the mountains is biased toward lower distance from the nearest road (<u>*x*</u>_*obs*_: 409.6 m, <u>*x*</u>_*sim*_: 1616.0 m, *P* < 0.001; Table 5). The median sampling distance from roads of historical records is a mere 55 m, compared to the simulated median sampling distance of 1,089 m (Figure 4C). While significant differences were detected between the historical and recent sampling data, both groups are similarly biased from random sampling (Fig. 4 D-F, Table 5), pointing to the difficulty of sampling mammals in the mid-elevation areas with steep relief and limited road access.

**Fig. 4.**
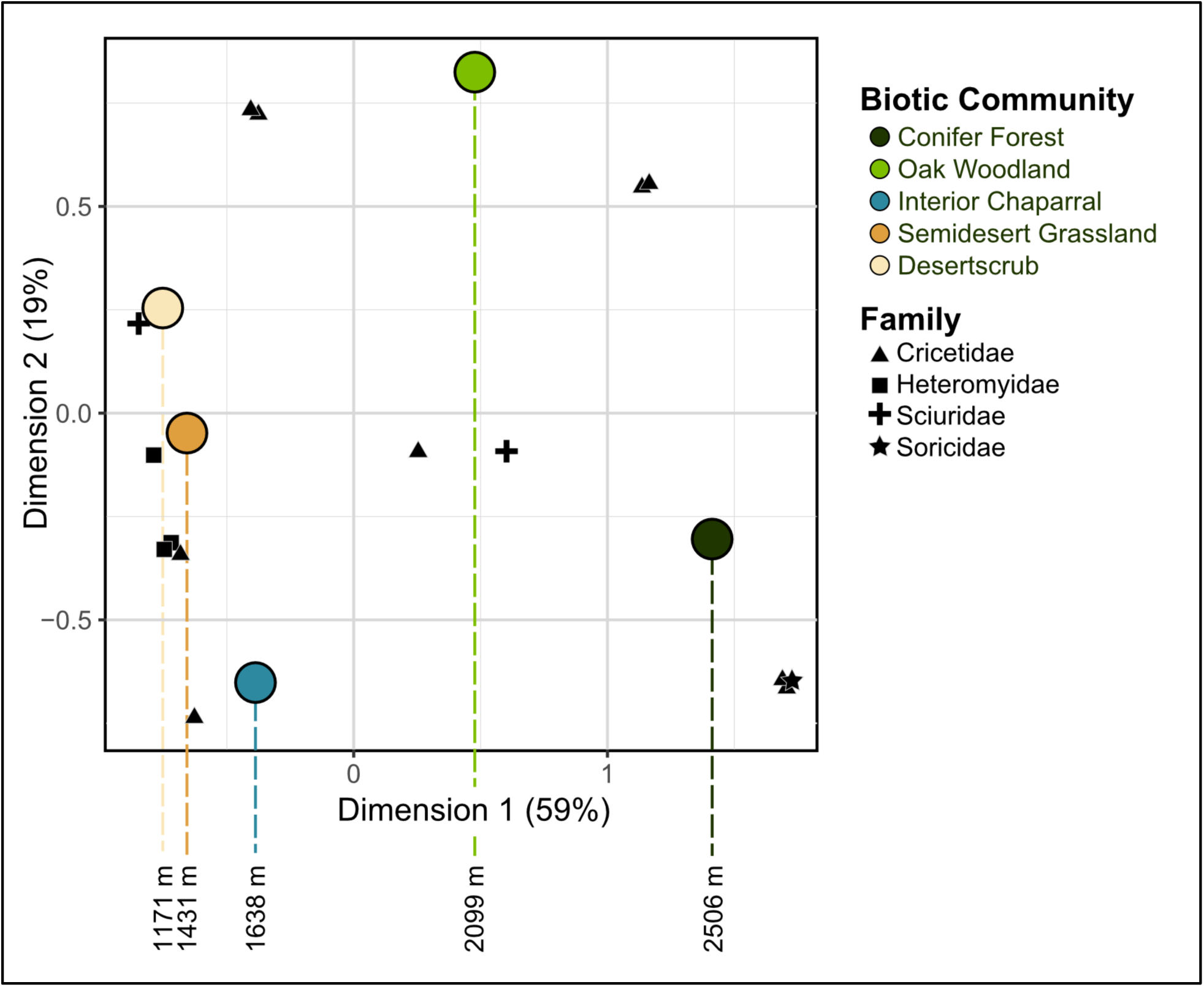
Community comparison across elevational biomes. Principal coordinate analysis (PCoA) of presence-absence data for the five biotic community types sampled in our collections. To highlight the strong elevational component of Dimension 1 (lower elevations on the negative end, higher elevations on the positive end), the mean elevation of each biotic community is listed below the plot.

Our 2021-2023 survey extends the known elevational ranges for 8 of 15 species we sampled compared to the historical dataset based on minimum and maximum sampled elevation, including 1 species with a downslope extension (*Reithrodontomys megalotis*, -320 m), 6 species with upslope extensions (ranging from +11 m to +394 m) and 1 species with both upslope and downslope extensions (*Sigmodon ochrognathus*; Table 4). The historical sample size per species ranged from 257 records (*P. boylii*) to only a single historical record (*R. megalotis*), indicating variation in the statistical confidence for the historical elevational ranges of these species. However, considering the 5 species having both ≥15 historical records from the Santa Catalinas and ≥15 records in our survey, all 5 species had upslope range extensions, including 2 species with extensions over +100 m (*Chaetodipus intermedius* and *Neotoma albigula*; Table 4). Thus, we have higher confidence in the elevational extensions for 5 of these 8 species, although insufficient presence-only data from historical and recent sampling to differentiate whether these new occurrences are elevational range extensions (based on statistical sampling) or actual range expansions (based on actual animal movement upslope).

**Table 4.** Comparison of minimum and maximum elevations (in meters, m) between historical and recent occurrences for which our 2021-2023 field survey extended elevational distributions. Range extensions marked with asterisks (*) are considered to have higher confidence given that their historical and modern sample sizes are each ≥15 individuals.

|  | Minimum elevation (m) |  |  | Maximum elevation (m) |  |  |
| --- | --- | --- | --- | --- | --- | --- |
| Species (*n* this study; *n* total ) | Historical | Recent | Extension | Historical | Recent | Extension |
| *Chaetodipus baileyi* (25; 243) | 845 | 1167 |  | 1432 | 1480 | +48* |
| *Chaetodipus intermedius* (36; 51) | 903 | 1149 |  | 1332 | 1538 | +206* |
| *Neotoma albigula* (17; 72) | 838 | 1148 |  | 1968 | 2088 | +120* |
| *Neotoma mexicana* (20; 42) | 1922 | 2065 |  | 2742 | 2791 | +49* |
| *Peromyscus boylii* (112; 369) | 865 | 1325 |  | 2413 | 2424 | +11* |
| *Reithrodontomys megalotis* (17; 18) | 2384 | 2064 | -321 | 2384 | 2423 | +39 |
| *Sigmodon ochrognathus* (9; 12) | 1383 | 1174 | -210 | 2171 | 2094 |  |
| *Sorex monticolus* (1; 8) | 2339 | 2789 |  | 2395 | 2789 | +394 |

### Comparison of co-occurring surveys of small mammals, plants, and arthropods

To verify that the terrestrial small mammal communities we sampled from 2021-2023 indeed correspond to different biomes in the Santa Catalina Mountains, we compared their species diversity with co-occurring communities of plants and ground-dwelling arthropods that were sampled by Moore et al. (2013) and Meyer et al. (2015), respectively. We find a strong positive relationship between the minimum species richness of all plants and small mammals across our 10 sampled sites (Fig. 5A; *r* = 0.710; *P* = 0.021). This pattern is explained somewhat more by uncommon plants (*r* = 0.716; *P* = 0.020) than common plants (*r* = 0.591; *P* = 0.072), as defined by Moore et al. (2013), even though fewer species per site were typically present (Table 6). Similarly, the minimum species richness of all arthropods is positively related to that of small mammals (Fig. 5B; *r* = 0.790; *P* = 0.007), a pattern that is driven mainly by the speciose Coleoptera (*r* = 0.690; *P* = 0.027) and Araneae (*r* = 0.790; *P* = 0.007; Table 6). The highest species richness was found at the SGN site (Semidesert Grassland north) for plants, arthropods, and small mammals (n=131, n=128, and n=8 respectively; Table 6).

**Fig. 5.**
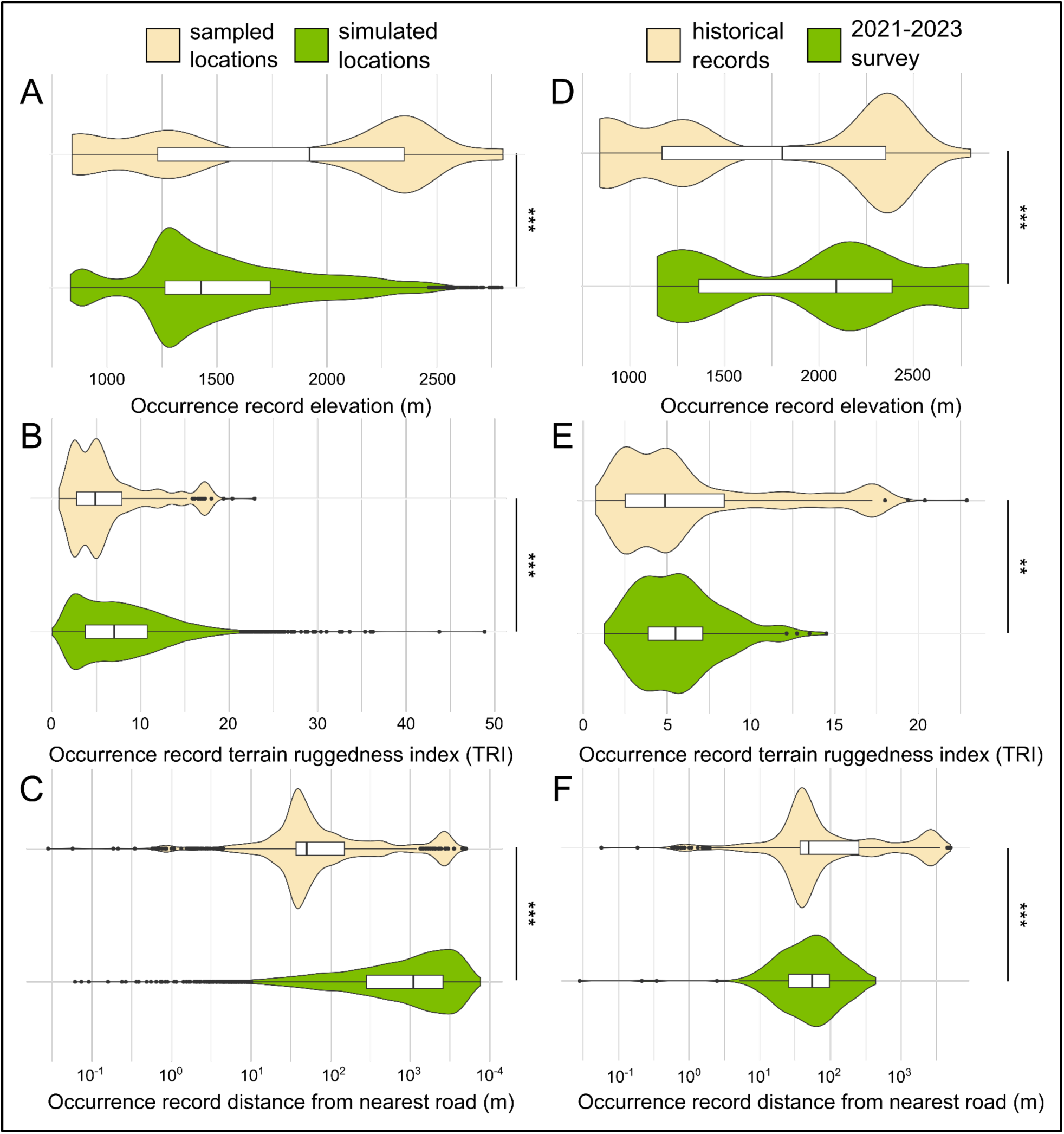
Box-and-violin plots of observed (*n* = 1,964) and simulated (*n* = 10,000) (A-C) and historical (*n* = 1600) and newly-sampled (*n* = 364) (D-F) mammal occurrences from the Santa Catalina Mountains with respect to A) elevation, B) terrain ruggedness index, and C) distance from nearest road. Asterisks represent significant differences of univariate components of MANOVA with ** indicating *p* < 0.01 and *** indicating *p* < 0.001 (See also Table 5).

**Table 5.**
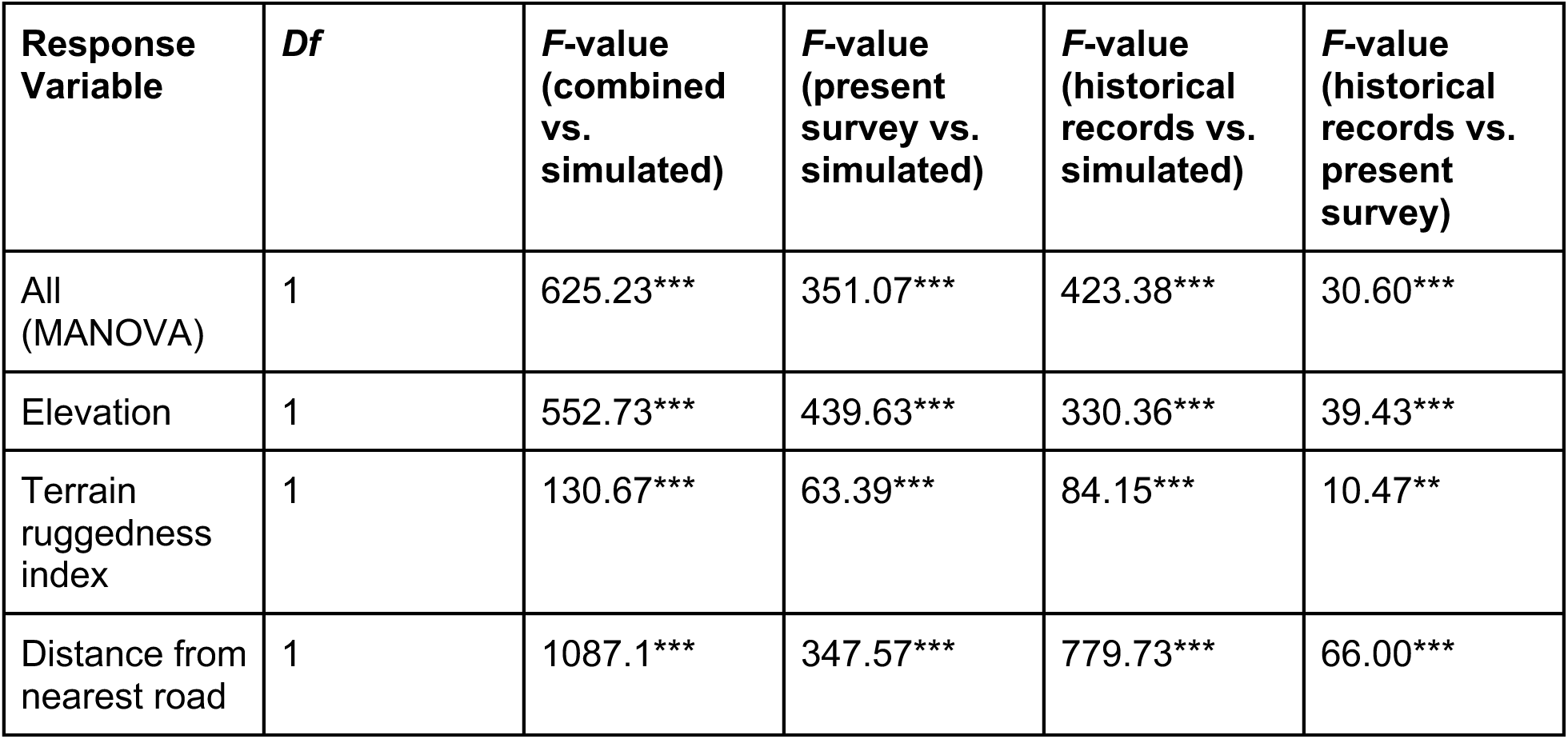
Summary table for MANOVA and univariate ANOVAs of sampling effort bias analyses of all filtered historical records and unique records from the current survey. Asterisks indicate level of significance: ** indicates *p* < 0.01, *** indicates *p* < 0.001.

**Table 6.** Comparison of species richness per biotic community for plants (Moore et al. 2013) and arthropods (Meyer et al. 2015) relative to the terrestrial small mammals surveyed in this study along the same transect of the Catalina Highway. Elevational midpoint is given from sites sampled in our survey.

|  |  |  |  | Species richness |  |  |  |  |  |  |  |  |
| --- | --- | --- | --- | --- | --- | --- | --- | --- | --- | --- | --- | --- |
| Site | Biotic Community | Slope | Elevation (midpoint) | Small mammals | Plants all | Plants common | Plants uncommon | Arthropods all | Orthoptera | Myriapoda | Coleoptera | Araneae |
| SDS | Desertscrub | S | 1180 | 6 | 74 | 44 | 30 | 85 | 5 | 5 | 31 | 44 |
| SGS | Semidesert Grassland | S | 1499 | 5 | 88 | 52 | 36 | 97 | 6 | 7 | 35 | 49 |
| ICS | Interior Chaparral | S | 1907 | 2 | 27 | 10 | 17 | 36 | 4 | 6 | 10 | 16 |
| OWS | Oak Woodland | S | 2130 | 3 | 50 | 26 | 24 | 77 | 4 | 7 | 27 | 39 |
| CFS | Conifer Forest | S | 2397 | 5 | 45 | 25 | 20 | 75 | 3 | 7 | 27 | 38 |
| CFP | Conifer Forest | Peak | 2776 | 4 | 43 | 24 | 19 | 76 | 4 | 6 | 36 | 30 |
| CFN | Conifer Forest | N | 2293 | 3 | 23 | 10 | 13 | 54 | 5 | 2 | 25 | 22 |
| OWN | Oak Woodland | N | 2075 | 5 | 30 | 9 | 21 | 87 | 4 | 5 | 39 | 39 |
| ICN | Interior Chaparral | N | 1329 | 4 | 54 | 32 | 22 | 84 | 5 | 4 | 35 | 40 |
| SGN | Semidesert Grassland | N | 1358 | 8 | 131 | 61 | 70 | 128 | 15 | 6 | 48 | 59 |

## Discussion

### Modern, holistic surveys are needed to understand system-wide sky island trends

To our knowledge, this study represents the most complete attempt to assess the diversity of small mammals in the Santa Catalina Mountains since Lange’s (1960) review 65 years ago. Our survey was guided by principles of holistic preservation (Galbreath *et al*. 2019; Thompson *et al*. 2021) and the extended specimen (Lendemer *et al*. 2020; Hardisty *et al*. 2022), with goals of systematically preserving multiple tissues from each sampled animal, cataloging whole organism vouchers for disparate future studies, and making these data easily accessible online. Due to permitting restrictions that limited our lethal vouchering to 3 animals per species per site, we also aimed to maximize the collection of non-lethal data in the form of ear clips, feces, and ectoparasites when available (Fig. 2), plus photographs and measurements from all animals, to help future workers verify our species identifications as taxonomies change (e.g., 234 more species of mammals have been recognized in North America since 2005; (Burgin *et al*. 2025). As a result, our survey bridged a ∼40-year lull in small mammal specimen preservation from the Santa Catalina Mountains (Fig. 1), extending the historical baseline of occurrence records while preserving frozen tissues from many localities for the first time.

Our survey highlights the need for additional surveys using the same holistic approach on other Madrean Sky Islands. For example, a recent US Forest Service camera-trap survey on the Rincon Mountains did not report shrews or rodents other than sciurids, underscoring the difficulty identifying small nocturnal mammals from images alone (i.e., without measuring traits like hind foot length, total length, and tail length; Swann and Perkins 2013). The last comprehensive mammal surveys for other Madrean Sky Islands in Arizona took place between the 1930s and 1950s, chiefly on the Chiricahua, Pinaleño, and Huachuca Mountains (Cahalane, 1939; Hoffmeister, 1956; Hoffmeister & Goodpaster, 1954). These surveys included collecting new specimens, compiling data on existing specimens, and/or listing accounts of local observations. However, most other sky islands in the Madrean system remain undersampled or entirely unsampled, especially those in Mexico (see Rivera and Upham, this volume). More recently, western New Mexico’s sky islands have received dedicated attention, including surveys of the Animas Mountains in the 1980s (Cook 1986) and the Gila Mountains from 2012-2020 (part of the southern Colorado Plateau rather than Madrean Sky Islands; Jones *et al*. 2021). The latter survey included camera trapping, small mammal collecting, and compiling historical records, yielding 2,919 new voucher specimens and preserving thousands of frozen tissues that have enabled downstream work (e.g., Malaney *et al*. 2023; Androski *et al*. 2025).

We emphasize that preserving frozen tissues using modern techniques such as liquid nitrogen and DNA/RNA shield dramatically increases the value of each voucher specimen for future investigations, a position we share with other researchers (McLean *et al*. 2016; Dunnum *et al*. 2017; Galbreath *et al*. 2019; Greiman *et al*. 2020; Thompson *et al*. 2021). Preserved tissues enable key downstream uses including: (i) characterizing microbial symbionts that inhabit small mammals (e.g., viruses, fungi, bacteria; Weinstein *et al*. 2021; Salazar-Hamm *et al*. 2022; Chen *et al*. 2023; Zhang *et al*. 2025), which can reveal differential tropism for modeling cross-species transmission risk (Cook *et al*. 2020; Carlson *et al*. 2022; Colella *et al*. 2023; Paul *et al*. 2025); (ii) using gene expression to measure host immunological responses to potential pathogens, which may show host disease tolerance consistent with co-evolutionary dynamics (Brook *et al*. 2023; Foley *et al*. 2024; Morales *et al*. 2025); and (iii) developing genomic and pan-genomic resources, which can reveal novel aspects of evolution and adaptation in small mammals (Callez *et al*. 2025; Fang and Edwards 2025) yet requires vouchering to ensure long-term data durability (Buckner *et al*. 2021). Generating genomic data from multiple trophic levels, spanning from microbial symbionts to vertebrate hosts and the diversity of nearby arthropods and plants (e.g., Fig. 6), will help illuminate how the interconnected parts of the Madrean Sky Islands ecosystem are responding to larger-scale change. In this way, we view the holistic sampling of one survey as enriching other surveys on other mountains, particularly when these data are digitally linked together for queries by site, specimen, species, or publication (Fawcett *et al*. 2022; Hardisty *et al*. 2022). Thus, a vision for region-wide comparative analyses of mammal ecological and evolutionary dynamics is within reach.

**Fig. 6.**
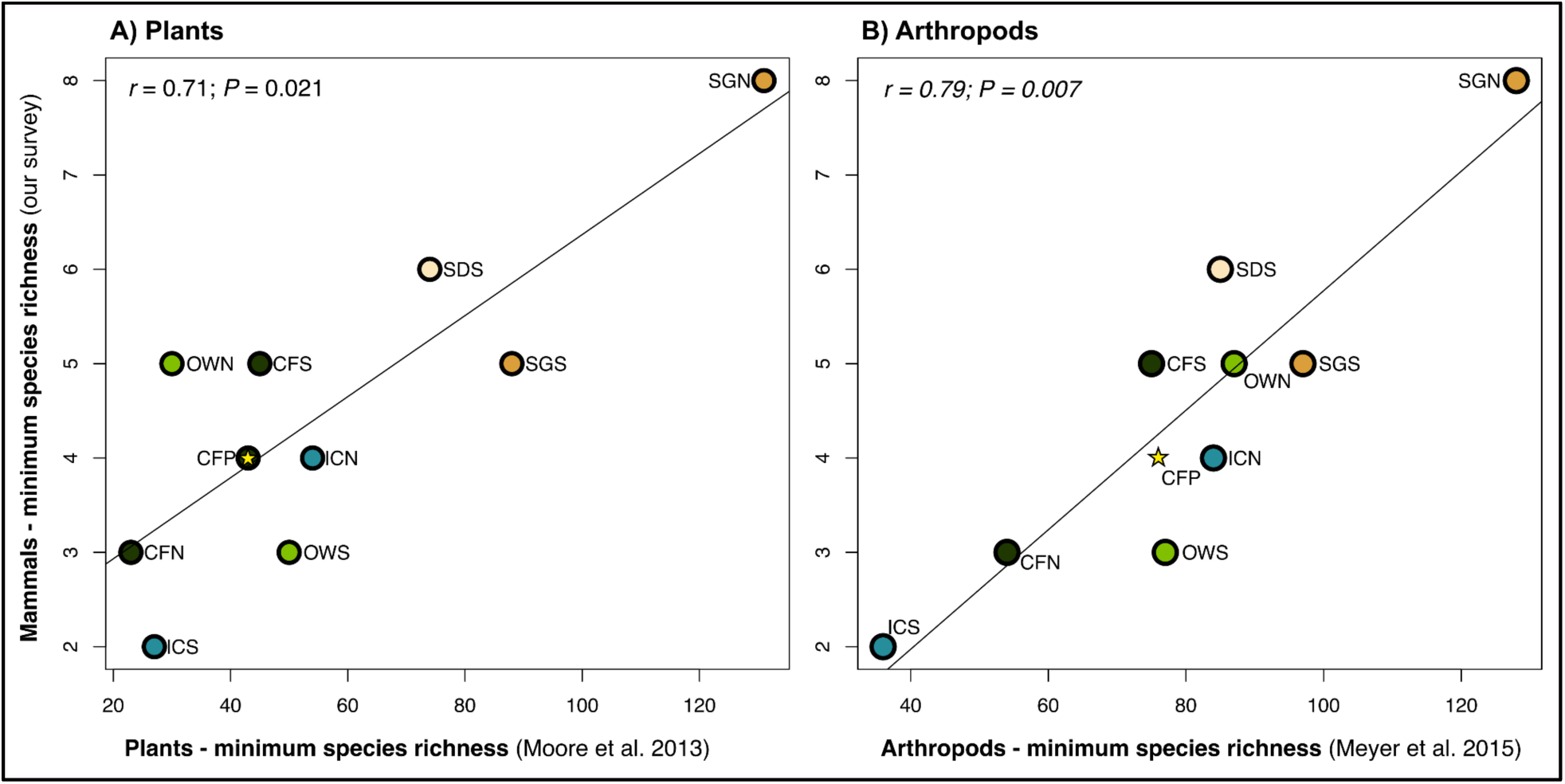
Comparison of species richness of terrestrial small mammal communities surveyed in the Santa Catalina Mountains for this study (10 sites, Nov 2021–Jul 2023) relative to surveys of other taxa in the same biomes: (A) plants, sampled in Aug 2011 (Moore et al. 2013); and (B) ground-dwelling arthropods (Orthoptera, Myriapoda, Coleoptera, and Araneae), sampled in May and Sep 2011 (Meyer et al. 2015). Values are representative of minimum species richness in a biome (rather than actual), given that transect-based sampling designs were used for all taxa. Biome abbreviations follow those given in Table 1 and colors follow Fig. 1A.

Although some view the lethal vouchering of wildlife as an anachronism from an earlier era of zoological research (Pape 2016; Waeber *et al*. 2017; Byrne 2023), the approach to holistic specimen preservation described here enables singular insights to a wide range of cutting-edge research questions that alternative sampling methods cannot address (Ceríaco *et al*. 2016; Malaney and Cook 2018; Nachman *et al*. 2023; Paul *et al*. 2025). Nevertheless, working with natural resource managers to alleviate potential concerns about vouchering is needed. We suggest that pairing lethal vouchering with non-lethal sampling of vouchered populations (as performed here; Fig. 2) can represent a kind of middle path for future Madrean Sky Island surveys: protecting sensitive taxa with reasonable lethal limits while systematically vouchering populations across elevational gradients. Doing so will build a rigorous time-series of holistically preserved specimens to add to the historical baseline of traditionally preserved (typically skin and skull) specimens across the region. Critically, population-genetic thresholds suggest that n=25-30 animals per species per timepoint are needed to reasonably approximate the levels of allelic diversity in most populations, as established based on tests of birds, small mammals, and insects (Hale *et al*. 2012) as well as sheep (Flesch *et al*. 2018). If those target species are small mammals, then using non-lethal sampling to help reach that threshold is reasonable; however, if the goal is to examine microbial symbionts of tissues such as lung, liver, and gut, then greater lethal sampling is needed. Ultimately, demonstrating the value of holistic vouchering practices is a shared responsibility that can be helped forward in multiple ways, including regional gap analyses identifying missing biodiversity data and integrative studies that emphasize the interconnected ecosystem within each animal.

### Modern surveys can extend known elevations of mammal species

Based on the findings of surveys of plants and arthropods in the Santa Catalina Mountains, we predicted that our modern sampling of terrestrial small mammals would show patterns consistent with species shifting their elevational distributions upslope compared to historical surveys. Our survey illustrates that over half of the species sampled (8/15) exhibit extra-limital occurrences relative to the 140-year historical baseline on this mountain (Fig. 1; Fig. 2; Table 4). While defining elevational ranges carries inherent biases of the sampling process (see Fig. 5; reviewed in Linck 2025), we find that 5 species have well-supported elevational changes (n ≥ 15 historical and recent observations; asterisks in Table 4). All 5 species exhibited upslope extensions — including +206 m for the *Chaetodipus intermedius* and +120 m for the *Neotoma albigula* — consistent with other surveys where montane species are moving upslope, including in the Santa Catalina Mountains (Rowe *et al*. 2010; Brusca *et al*. 2013; McCain *et al*. 2021; Holzmann *et al*. 2023). However, we also note that the historical sampling of any Madrean Sky Island is unlikely to be sufficient to detect true range expansions (see Upham and Rivera, this volume), particularly since historical sampling effort is generally unknown. As a result, the elevational changes of these 8 species must be considered range *extensions* until future survey work is conducted (Frey 2009). The preponderance of upslope elevational extensions we recorded is nevertheless interesting. We suggest that future studies with a focus on spatial occupancy would be helpful in determining whether these extensions are actually shifts or expansions that are occurring in response to changes in their abiotic environment and available biotic resources.

### Small mammal species richness in the Santa Catalinas mirrors that of other taxa

Elevation is the primary signal in our analysis of small mammal species composition over biotic communities (Fig. 4). Part of the impetus for our study stemmed from determining how small mammal species richness differs across biotic communities and relative to other taxa that have been surveyed in the Santa Catalina Mountains, namely plants and ground-dwelling arthropods. Our survey of small mammals indeed supports this prediction, in terms of species clustering with biotic communities (Fig. 4). Plant surveys conducted between 1963 and 2011 reported upslope shifts for 23 plant species, including shifts of over 250 m for the lower elevations of alligator juniper (*Juniperus deppeana* var. *deppeana*) and Gambel oak (*Quercus gambelii*; Whittaker and Niering 1964, Brusca et al. 2013). Arthropod communities similarly appear to be moving upslope, as supported by the tight clustering of species within plant-defined biotic communities (Meyer et al. 2015). However, the 2011 survey of arthropods did not identify explicit elevational shifts due to the paucity of historical Santa Catalinas sampling for many species (Moore et al. 2013, Meyer et al. 2015). These authors found the highest species richness of plants and arthropods in low- to mid-elevation Semidesert Grassland (there called Desert Grassland and Oak-Grassland) and Desertscrub sites, and the lowest diversity in Conifer Forest (called Pine Forest; Moore et al. 2013; Meyer et al. 2015). We similarly recorded the highest minimum species richness in Semidesert Grassland (9 total species; 8 on the north slope, 5 on the south slope), but had 3 biotic communities tied for the lowest minimum small mammal diversity (6 total species; see Table 2, Table 6). Ordination by habitat similarity shows a clear elevational signal in mammals, plants, and arthropods (Fig. 4 in this study as compared to fig. 4 in Moore et al. 2013, fig. 2 in Meyer et al. 2015). Small mammal communities are most diverse at the same sites where plant diversity is highest (Fig. 6), supporting the expectation that higher trophic levels are ecologically dependent on lower ones (see fig. 10 in Rosenzweig and Winakur 1969).

### Surveys in accessible mountains still reveal new insights on mammalian species

Our survey of the Santa Catalina Mountains contributed 5 noteworthy observations that highlight the need for continued sampling of even the most road-accessible sky islands. First, we collected the first known specimen of *Reithrodontomys fulvescens* from the Santa Catalina Mountains: [ASUMAC009016](https://cvcoll.org/portal/collections/individual/index.php?occid=1093034&clid=0), a subadult male with semi-scrotal testes collected on 3 July 2023 from Bear Wallow near the General Hitchcock Rd. overpass, 32.4231380 °N, 110.7339500 °W. The animal was trapped in moist riparian habitat downstream from the capped Bear Wallow spring and discriminated from *R. megalotis* and *R. montanus* based on a combination of cranial, skeletal, and external morphological features: tail length > 80 mm (91 mm), baculum length > 7 mm (7.61 mm), and the presence of a lingual fold in upper M3 that gives the molar an ‘E’ shape (as opposed to the C-shaped molars of *R. megalotis* and *R. montanus*; Hoffmeister 1986). Our searches of publicly available digital records indicate that the nearest records of this species consist of two animals (UAZ 26887, 26895) collected approximately 32 kilometers by air to the south, on the bank of Rincon Creek within riparian habitat of what appears to be a Desertscrub biotic community. The fact that these two localities differ so greatly in elevation and other habitat characteristics suggests that *R. fulvescens* may be associated more closely with riparian zones than any particular biotic community, as was noted in the Animas Mountains (Cook 1986). Other authors note that this species prefers habitat with multilayered herbaceous cover, which is consistent with the habitat where we recorded this individual (Spencer and Cameron 1982; Clark *et al*. 1998). We interpret this new detection as a range extension into the Santa Catalinas, given the lack of background occurrences of this species from the mountain and region (Frey 2009).

Second, on 2 July 2023 we detected individuals of both *Neotoma albigula* (1 subadult male, 1 lactating female) and *Neotoma mexicana* (1 juvenile female) in an apparent contact zone of the species in a north-slope Oak Woodland community that had been severely impacted by the Bighorn Fire of 2020. This site contained numerous burned snags of pine, oak, and juniper, and amid the opening of the canopy, grasses had been able to proliferate. We speculate that the fundamental change in this biotic community may have allowed *N. albigula* to move into communities previously more favorable to *N. mexicana*. Efforts are underway to better understand the niche partitioning of *N. mexicana* and *N. albigula* in the Santa Catalinas, including relative to dietary and gut microbial axes (G. De Leon, pers. comm., 18 Sept 2025).

Third, we did not encounter *Sigmodon arizonae* in our study as expected given the habitats we sampled. However, we did encounter its congener *Sigmodon ochrognathus* in Semidesert Grassland and the burned north-slope Oak Woodland communities. Those latter observations substantially increased the areas and elevations from which this species had previously been recorded in the Santa Catalina Mountains (Table 4). Considering the limited number of representatives of this species in the historical dataset, we can conclude that this is a range extension of the species in the Santa Catalina Mountains (Frey 2009). However, Davis & Dunford (1987) argue that despite the ease of accessibility of the Santa Catalina Mountains for collectors since the early 20th century, the first specimen of *S. ochrognathus* was not collected until 1967, and argue that this historical background of collecting is substantial enough to provide evidence to support the hypothesis that *Sigmodon ochrognathus* is exhibiting a range expansion across the broader Madrean Sky Island region. The authors suggest this species is dispersing northward and westward independent of human involvement, to the point where it is now relatively common throughout the region (Davis and Dunford 1987). However, within the Santa Catalinas, it remains an open question whether *Sigmodon ochrognathus* is experiencing a trend of range expansion. It is worth exploring whether the proliferation of Semidesert Grassland communities at middle and high elevations due to recent wildfires may have benefited *S. ochrognathus* in particular, since, based on our historical dataset, *S. arizonae* appears to have been collected only within Desertscrub communities below 1000 m (i.e., lower than our lowest sampling locality; Davis and Dunford 1987).

Fourth, our survey also failed to recover a few species we reasonably expected to recover based on historical records, namely *Dipodomys merriami*, *Dipodomys spectabilis*, *Xerospermophilus tereticaudus*, and *Notiosorex crawfordi*. These species are principally low-elevation species associated more with desertscrub than other biotic communities (Hoffmeister 1986). We cannot conclude that these species are absent from the sites we sampled based on the inexhaustive nature of our survey, but we do note that historical records indicate a distribution overlapping at least two of our sites for *Dipodomys merriami, Xerospermophilus tereticaudus*, and *Notiosorex crawfordi* (Table 3). These species are distinct enough from those that were sampled that we find it unlikely that they could have been misidentified and released without vouchering. Three additional species of sciurids (*Otospermophilus variegatus*, *Sciurus aberti*, and *S. arizonensis*) and the two species of pocket gophers (*Megascapheus bottae*, *M. umbrinus*) constitute the remaining small mammals that we were unlikely to collect in our Sherman traps, although we did observe the presence of *Otospermophilus variegatus*, *Sciurus aberti*, and *Megascapheus* sp. in at least one of the sites where we trapped. The intentional introduction of *Sciurus aberti* to the Santa Catalinas in winter 1940 has long been pointed to as contributing to the decline of *Sciurus arizonensis* in the Santa Catalina Mountains (Lange 1960; Davis and Brown 1988).

Finally, the species boundary of the montane, short-tailed (< 80 mm) *Peromyscus* that occurs at high elevations (typically > 2,200 meters, but see some exceptions in Fig. 3) in the Santa Catalinas warrants further investigation, especially in the context of occurrences on other Madrean Sky Islands. We provisionally refer to this species as *Peromyscus melanotis* because of longstanding evidence of its occurrence at high elevations and a lack of evidence of the existence of other short-tailed *Peromyscus* from the mountains (Bowers *et al*. 1973; Bowers 1974; Natarajan *et al*. 2015). However, the appropriate name(s) to apply to these short-tailed *Peromyscus* is mired in historical inconsistency. The subspecific name *rufinus* Osgood, 1909 was used by Lange (1960) to refer to this high-elevation *Peromyscus* in the Santa Catalinas (“*P. maniculatus rufinus*” above Summerhaven, >2,280 m) while low-elevation *Peromyscus* was assigned to “*P. maniculatus sonoriensis*” (near Molino Basin). Our survey did not recover any short-tailed *Peromyscus* below 2,355 m, possibly because our lowest locality surveyed (1146 m) was too high to obtain the supposed lowland form. In our physical verification of historical specimens of low-elevation “short-tailed” *Peromyscus*, some apparently short-tailed *Peromyscus* were actually *P. eremicus* or *P. boylii* with bobbed tails (i.e., damaged), which could explain the three low-elevation records of ‘*P. melanotis*’ shown in Figure 3. However, we also note that these specimens were collected in a riparian area, habitat which could allow a high-elevation species to reach lower elevations. In either case, we suggest that the identity of historical records of *Peromyscus* identified with epithets of *maniculatus*, *sonoriensis*, *leucopus*, *melanotis*, and *rufinus* from the Santa Catalinas, especially those from low elevation, should be regarded with caution and confirmed with genetic and specific collection locality data whenever possible. Untangling the species boundaries of this group would ideally entail analyses of genome-wide polymorphisms and sampling from not only the Santa Catalinas, but other Madrean Sky Islands and the ‘mainlands’ of the Sierra Madre Occidental and Colorado Plateau to better understand which species occur in these areas (A. Fenlon, pers. comm., 24 Sept. 2025). These observations of terrestrial small mammals illustrate that new insights are still possible even for sky islands that are accessible by road and relatively frequently sampled, supporting the claim that none of the Madrean Sky Islands are completely sampled (Rivera and Upham, this issue).

### Sky islands are rapidly changing environments - are communities keeping up?

We recommend that future sampling of sky islands should also be conducted with an eye toward fire impacts and recovery. Our survey conducted sampling at two sites impacted by the Bighorn Fire of 2020, which burned 3.5% of the Santa Catalinas by area, much of which was montane woodland and forest (NIFC Authoritative Content 2023). Most information about fire impacts on small mammals of the Madrean Sky Islands has focused on the endangered Mount Graham Red Squirrel, which is endemic to the Pinaleño Mountains (Koprowski *et al*. 2006; Merrick *et al*. 2021). Future studies should aim to facilitate nuanced understanding of faunal dynamics in these unusually isolated sky island communities. In the Santa Catalinas, as in other montane regions, higher-elevation habitats include relatively high proportions of locally endemic taxa (Meyer *et al*. 2015) and may recover more slowly or not reach pre-burn levels of species richness due to the local extirpation of species. This risk becomes more likely as climate change forces high-elevation plant species further upslope (Brusca et al. 2013) and further fragments the already patchy sky-island habitats (Yanahan and Moore 2019). The region’s arid climate may also have slowed habitat recovery from fire damage (Lv *et al*. 2025), a hypothesis that could be tested via longitudinal studies of fire-impacted areas on the mountain. Doing so would represent a worthwhile extension of our survey to determine the timing and pattern of recolonization from the Bighorn Fire, including whether these patterns differ by elevation, fire severity, and biotic community (Culhane *et al*. 2022).

## Conclusion

Our investigation of mammal communities in the Santa Catalina Mountains provides a new baseline for future studies of montane biodiversity in the region and serves as a resource for future inquiry. Our study also confidently extends the known elevational range of 5 rodent species in our study upslope relative to the baseline of historical sampling, as well as provides the first known occurrence of another species. Our study begs the question as to whether the mammals of the Santa Catalinas are responding to a variety of abiotic and biotic changes: principally, the upslope dispersal of plants on which rodents depend, rising temperatures and erratic rainfall that has impacted plant distributions, and the spread of grassland following the 2020 Bighorn Fire. Zooming out, these kinds of changes are likely common across other mountains of the Madrean Sky Islands. The direct effects of climate change (e.g., longer-term increases in temperature and more frequent and severe droughts) are predicted to worsen by 2050, thereby exacerbating indirect effects on montane habitats (e.g., increasing risk and severity of wildfire damage as a result of changing temperature; Yanahan and Moore, 2019). We argue that there is a clear need for the study of montane biodiversity in the Madrean Sky Islands. Small mammal surveys like those our team conducted, especially when integrated with historical collecting efforts, contribute to a growing knowledgebase for understanding and mitigating the impacts of continued climatic changes. The historical baseline of specimen collections in the Santa Catalinas, which extends back to the 1880s, is now joined with a pulse of observations from our re-survey efforts 140 years later, enabling the key temporal comparison for measuring change. That simple “before and after” idea is indeed one of the most compelling reasons for 21st century mammalogy, and has motivated some of its most impactful insights (e.g., Tingley & Beissinger, 2009; Smith et al., 2013; Riddell et al., 2021). The voucher specimens produced by surveys such as ours are far more than a snapshot of the biota, but a “moving picture” for assessing the extent and timing of biotic change. Our results show that there is still much to be uncovered even from accessible mountains like the Santa Catalinas. In the context of continued global change, mammalogists have a crucial role to play as both documentarians of biotic change and advocates for a future in which biotic communities are healthy actors despite dramatic shifts in climate.

## Acknowledgments

We would like to thank the following individuals for assistance with permitting needed to perform this study: Neil Dutt and Tavia Carlson, Coronado National Forest; Christina Kondrat-Smith, AZGFD. We are indebted to Gilma De Leon, Edward Gilbert, Aven Gilbert-Kleker, Eduardo Gutierrez, Sharon Hall, Dorea Kleker, Rosie Liao, Kory Luedke, Katelin Pearson, Gregory Post, and Michael Vidaure for assistance in the field. We are grateful to the following museum staff who provided information on confirming IDs of historical specimens: Jacob Esselstyn (Louisiana State University Museum of Zoology); Nepsis Garcia, Cody Thompson (University of Michigan Museum of Zoology); Angela Hornsby, Sharon Jansa (Bell Museum of Natural History); Mark Omura (Harvard Museum of Comparative Zoology); and Thomas Trombone (American Museum of Natural History). We thank members of the ASU Natural History Collection and NEON Biorepository for general logistical support, especially Nico M. Franz. We provide special thanks to members of the Upham Lab of Mammal Phylogenetic Ecology for helpful comments, especially Alexander Fenlon, Ryan Nguyen, Gilma De Leon, Morgan Pierce, Julia Nitschmann, and Jessica Neumaier. Amanda Grunwald also provided helpful discussion of sky islands. Funding was provided by Arizona State University and grants from the National Institutes of Health (1R21AI164268 and 1R35GM156919 to NSU).

## Supplementary Data

Supplementary Data can be found at the publisher’s website.

**Supplementary Data SD1.** Listing of rodents, bats, and shrews given in Lange (1960), the last systematic survey of small mammals of the Santa Catalina Mountains before this study, as compared to the taxonomy of the Mammal Diversity Database v2.0 (MDD 2025). Species names in the MDD are bolded if they have changed in genus or species allocation since Lange (1960).

**Supplementary Data SD2.** Linear regressions of DEM elevation against verbatim elevation. Records are plotted in columns based on whether they exhibited residuals that fell within (left column) or exceeded (right column) a threshold standard deviation from the mean residual value: A) 1 SD, B) 2 SD, C) 3 SD. Records with residuals exceeding 3 standard deviations from the mean were excluded from subsequent analyses.

**Supplementary Data SD3.** Presence-absence matrix for Santa Catalinas habitats, based on recent sampling only.

